# Benefit Transfer Loops Turn Cheating into a Scaffold for Microbial Diversity

**DOI:** 10.1101/2023.11.21.568182

**Authors:** Jiqi Shao, Yinxiang Li, Shaohua Gu, Xiaoyi Zhang, Shaopeng Wang, Xueming Liu, Zhiyuan Li

## Abstract

Niche construction drives ecological dynamics, yet the tragedy of the commons predicts that non-contributing cheaters will undermine cooperation. Here, we studied microbial iron competition by combining dynamic modeling with benefit flow graphs, demonstrating that moderate cheating is not merely tolerated but essential for diversity. In small communities, mutual exploitation forms closed loops enabling steady or dynamic coexistence. In larger communities, we uncovered a paradox: increasing cheating breadth promotes community-level extinction, yet fosters higher biodiversity in surviving communities. We resolve this paradox by mapping ecological dynamics onto the topology of the “Maximal Benefit Transfer Graph”, which predicts community fate through its core structure. Broad cheating eliminates the self-loop core that drives competitive exclusion, but increases “terminator” sinks that cause collapse. However, when communities avoid these sinks, cheating aggregates the network and generates cyclic loops to enable coexistence. Thus, structured exploitation acts not as destabilizing vulnerability but as necessary architecture for biodiversity.

**Graphical Abstract:** 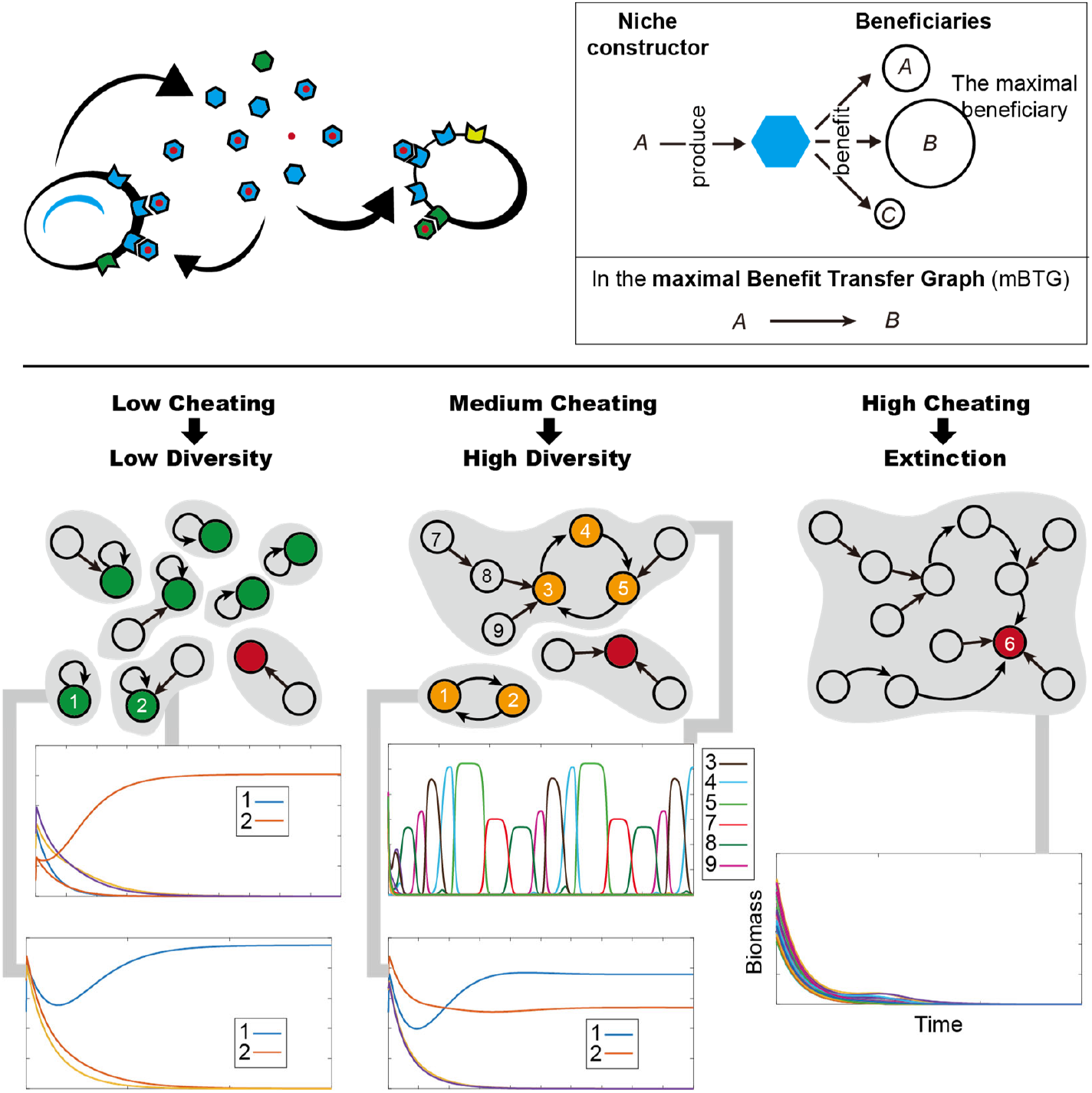

How does ‘cheating’ affect microbial biodiversity? By mapping the strongest benefit flows between species, we discovered a topological rule for survival. While too much cheating creates dead-ends that crash the system, moderate cheating connects species into self-sustaining loops. These “exploitation cycles” act as a scaffold, supporting high diversity and complex population.

## Introduction

Niche construction drives ecological dynamics[1], yet the “tragedy of the commons” predicts that non-contributing cheaters should undermine cooperative systems[2]. Microbial siderophores represent a classic model of this dilemma: microbes secrete costly iron-scavenging molecules as public goods, which are then uptaken via specific membrane receptors[3]. While functionally straightforward, siderophore systems display remarkable complexity stemming from two key features[4]: (1) extraordinary chemical diversity comprising at least a thousand structures[5]; and (2) high receptor specificity, creating “lock-and-key” patterns where receptors bind only specific siderophore subsets[6]. This specificity generates a rich ecological landscape where microbes possessing matching receptors can exploit “foreign” siderophores[7], forging intricate networks of competition, cooperation, and cheating[8].

This complexity makes siderophore-mediated interactions an ideal system for addressing a central puzzle in ecological theory: how does biodiversity persist in the face of cheating[9-11] ? Classical theory suggests cheaters should outcompete producers, yet producers and cheaters coexist broadly in nature [12]. Observations in the *Pseudomonas* genus further complicate this picture: while pathogenic strains often exhibit extreme strategies with isolated iron-interaction networks, more diverse environmental isolates likely to adopt “partial-producers” strategies that producing siderophores while cheating on others[6]. A fundamental challenge lies in bridging scales: how do fine-grained molecular attributes, including structural diversity and receptor specificity, give rise to macro-scale ecological patterns? While previous studies have explored siderophore dynamics in biofilms or pairwise competitions[13, 14], they often overlook the emergent properties of networks formed by multiple siderophore types [15-17]. Understanding how biodiversity persists in complex communities requires bridging the gap between molecular-level interaction networks and emergent ecosystem dynamics[18].

Here, we develop a general theoretical framework that integrates consumer-resource dynamics with graph-theoretic representation of benefit flows, to systematically address these questions. Through simulations and analytical derivations, we prove that cheating is actually required for stable and oscillatory coexistence in small communities. In large communities, we uncover a paradox: increasing the breadth of cheating heightens the risk of community-level extinction, yet fosters higher biodiversity within surviving communities. By mapping ecological dynamics onto a “maximal Benefit Transfer Graph,” we identify the graph’s core structure as a powerful predictor of ecological outcomes, explaining cheating’s paradoxical role: Increased cheating pervasiveness connects species through network percolation, simultaneously expanding the “terminator” core driving extinction and the cyclic core enabling coexistence. Thus, our findings reveal a universal topological rule: structured exploitation acts not as a destabilizing vulnerability but as a necessary architecture for diversity. This principle extends beyond iron competition to broadly explain persistence in systems governed by directed benefit transfers, from extracellular enzyme hydrolysis to complex metabolic cross-feeding networks.

## Results

### A Generalized Framework for Siderophore-mediated Interactions

To link molecular specificity with ecological dynamics, we developed an integrated framework coupling dynamic modeling with network topology (Fig. 1, SI Appendix, Section 1). Our dynamic model extends classical chemostat equations by incorporating diverse siderophore-receptor pairs, where growth depends on both iron uptake and primary metabolism (Fig. 1A). The model incorporates two key biological constraints: (1) a metabolic trade-off (parameter *α*), where species *i* allocate limited resources between growth (*α*_*i0*_) and siderophore production (*α*_*ij*_ for siderophore type *j*, under the constrain 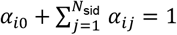); and (2) a receptor profile (parameter *v*), which dictates the specificity of iron uptake (*v*_*ij*_ represents the fraction of receptors dedicated to siderophore type *j* in species *i*, under the constrain 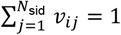).

**Figure 1.**
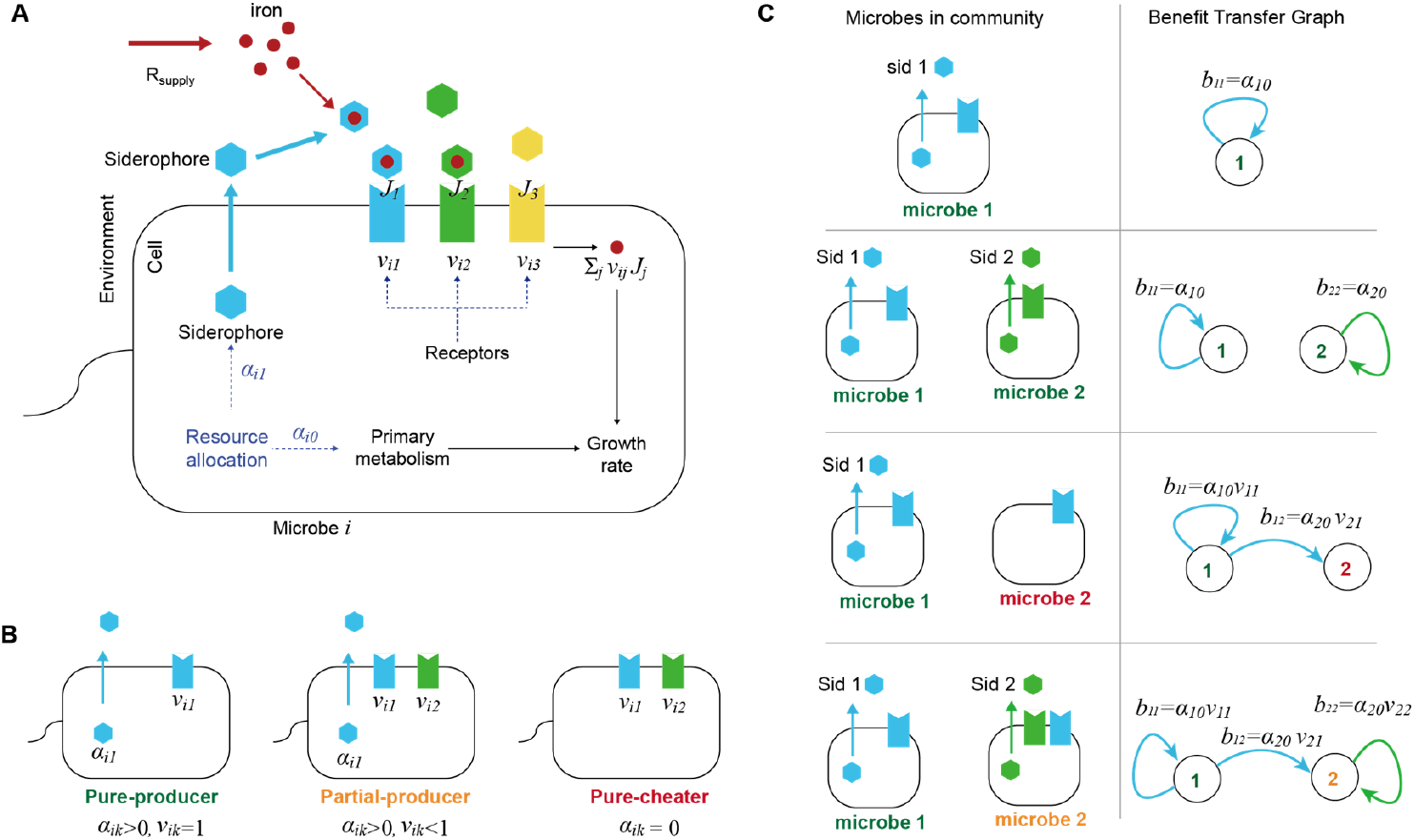
Framework for modeling siderophore-mediated interaction and benefit transfer in microbial communities. (A) Overview of the siderophore-mediated iron uptake. Microbes allocate internal resources between growth (*α*_*i0*_) and the production of siderophores (*α*_*ij*_ for *j* > 0). Secreted siderophores form siderophore-iron complexes, which are taken up via type-specific receptors with allocation fractions *v*_*ij*_. Different types of siderophores with their matching receptors are distinguished by colors. (B) Microbial iron-scavenging strategies. Microbes are categorized into three major classes by siderophore production and uptake patterns: (i) “Pure-producers,” which produce and exclusively utilize their own siderophores; (ii) “Partial-producers,” which produce/utilize their own siderophores and also exploit foreign siderophores; (iii) “Pure-cheaters,” which rely entirely on siderophores produced by others. (C) Benefit Transfer Graph (BTG). Left panels illustrate example siderophore-mediated interactions; right panels show their BTG representations: Nodes denote species, and directed edges represent benefit transfer from siderophore producers to beneficiaries. Edge colors correspond to siderophore types.

The dynamic model tracks three sets of variables: (1) Microbial biomass concentration *M*_*i*_ for each species *i* = 1,2, …, *N*_spe_ ; (2) concentrations of each siderophore type *R*_*j*_ (*j* = 1,2,…, *N*_sid_); (3) Free iron concentration *R*_iron_:

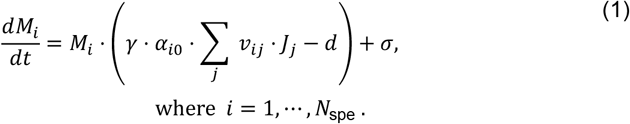

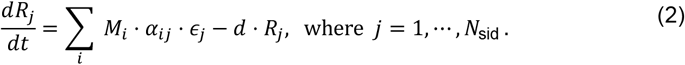

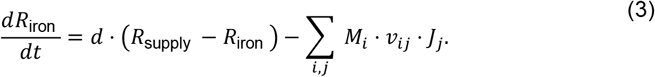

Varying parameters *α* and *v* generates a continuum of strategies, from “pure-producers” (exclusively utilizing their own siderophores) to “pure-cheaters” (no production) and “partial producers” (producing one siderophore while exploiting several other types) (Fig. 1B).

Crucially, we mapped these biochemical interactions onto a “Benefit Transfer Graph” (BTG) to quantify ecological dependencies (Fig. 1C). In this directed graph, nodes represent species, and edges represent the flow of growth benefits from a siderophore producer to its beneficiaries. An edge exists if a species possesses receptors matching another’s siderophore, with the weight *b*_*m*1,*m*2_ quantifying the specific growth gain species *m*2 derives from species *m*1’s production:

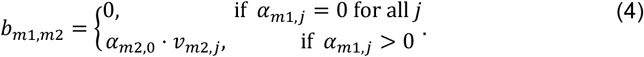

This abstraction allows us to translate complex kinetic systems into topological structures, analyzing how benefit transfers drive community assembly.

### Closed Benefit Loops Drive Transitions from Exclusion to Coexistence and Chaos

We first analyzed minimal community motifs to dissect the topological rules of coexistence. In single-species systems with only one producer (Fig. 2A), the Benefit Transfer Graph only consists of self-loops. Dynamic modeling reveals that siderophore-mediated positive feedback creates an Allee effect [19], leading to bistability where survival depends on initial biomass thresholds (Fig. 2B; SI Appendix, Section 2). Consequently, when two pure-producers compete, they are driven to competitive exclusion: either one species becomes completely dominant, or the system exhibits priority effects in which the final winner is determined by who is initially more abundant (SI Appendix, Section 3)

**Figure 2.**
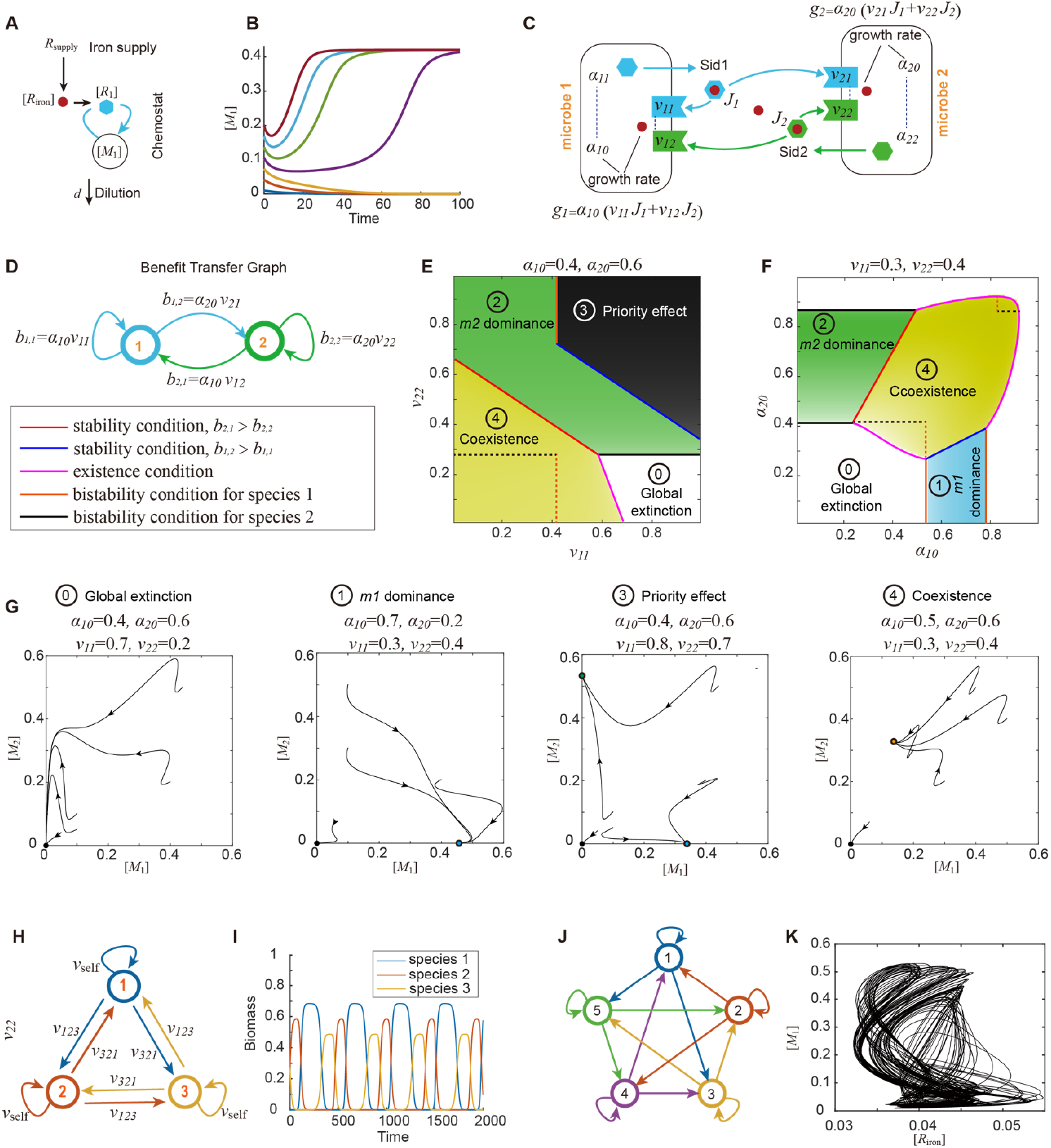
Rules governing siderophore-mediated iron interactions in single- and two-species models. A. Schematic of the single-species chemostat model with iron supply *R*_supply_ and dilution rate *d*. Variables [*R*_iron_], [*M*_1_], [*R*_1_] denote the concentration of free iron, microbial biomass, and siderophore, respectively. B. Time courses of biomass (*M*_1_) starting from different initial inoculations, illustrating the threshold-dependent survival. C. Schematic of a two-species system where two pure-producers compete. Each species secretes and exclusively utilizes its own siderophore type (blue for species 1, green for species 2). D. The Benefit Transfer Graph (BTG) corresponding to (C). Nodes represent species; edges represent the benefit flow. This graph features self-loops (*b*_1,1_, *b*_2,2_) and cross-species benefit transfers (*b*_1,2_, *b*_2,1_) E-F. Phase diagram spanning the receptor profile (*v*_11_-*v*_22_ for E) and growth allocation (*α*_10_ - *α*_20_ for F). Distinct ecological outcomes are color-coded and numbered. Color intensity is proportional to the total steady-state biomass. G. Representative state-space simulation dynamics projected onto the *M*_1_-*M*_2_ plane, parameters are from four different regimes in (E)-(F). Arrows denote the directionality of trajectories. Solid circles indicate stable fixed points. H. BTG of a three-species system. Blue, orange, and yellow arrows represent benefit transfers mediated by different siderophores produced by species 1, 2, and 3, respectively. Two rock-paper-scissors loops emerge: clockwise (characterized by *v*_321_) and counterclockwise (characterized by *v*_123_). Self-loops indicate self-utilization (*v*_self_) I. Representative time courses showing sustained oscillations in the system of (H). J. BTG of a five-species system. K. State-space projection onto the *R*_iron_-*M*_1_ plane showing a chaotic trajectory.

Stable coexistence emerges only when species adopt “partial-producer” strategies, forming a closed mutual cheating loop in the BTG (Fig. 2C–D). Analytically, we derived that each species must deliver greater growth benefits to its competitor than to itself (*b*_2,1_ > *b*_2,2_, *b*_1,2_ > *b*_1,1_, SI Appendix, Section 4). Strikingly, this mutual exploitation generates a synergistic rescue effect: in parameter regimes where single species would collapse, the combined siderophore pool allows the mutualistic pair to survive (Fig. 2E–G).

Expanding these loops to multi-species motifs identifies key topologies that enable dynamic coexistence. In three-species systems, an intransitive “rock–paper–scissors” BTG loop (Fig. 2H) generates sustained population oscillations (Fig. 2I). Bifurcation analysis demonstrates that increasing self-reliance drives a transition from oscillatory coexistence to competitive exclusion via Heteroclinic bifurcation[20], while balanced benefit flows favor stable coexistence over limit cycles (SI Appendix, Section 5).

As the interaction network grows more complex, such as in five-species motifs with overlapping loops (Fig. 2J), coupled feedback can drive the system into deterministic chaos (Fig. 2K). Similarly, increasing self-reliance progressively drives transition from chaotic dynamics to stable periodic cycles and, eventually, to exclusion. Collectively, these results demonstrate that closed benefit-transfer loops serve as the structural scaffold for diversity: short mutual loops support stable coexistence, while longer cycles enable dynamic fluctuations.

### The Paradox of Cheating in Large Communities

We extended our framework to complex ecosystems by simulating 1.5 · 10^6^ communities (*N*_spe_ = 50) with varying “cheating breadth” (the average number of foreign siderophores a species can exploit) and fractions of “pure-cheaters” (Fig. 3A). Simulation outcomes reveal an unintuitive paradox. On one hand, broad cheating and high pure-cheater ratio act as destabilizing forces, monotonically increasing the probability of community-wide extinction (Fig. 3B).

**Figure 3.**
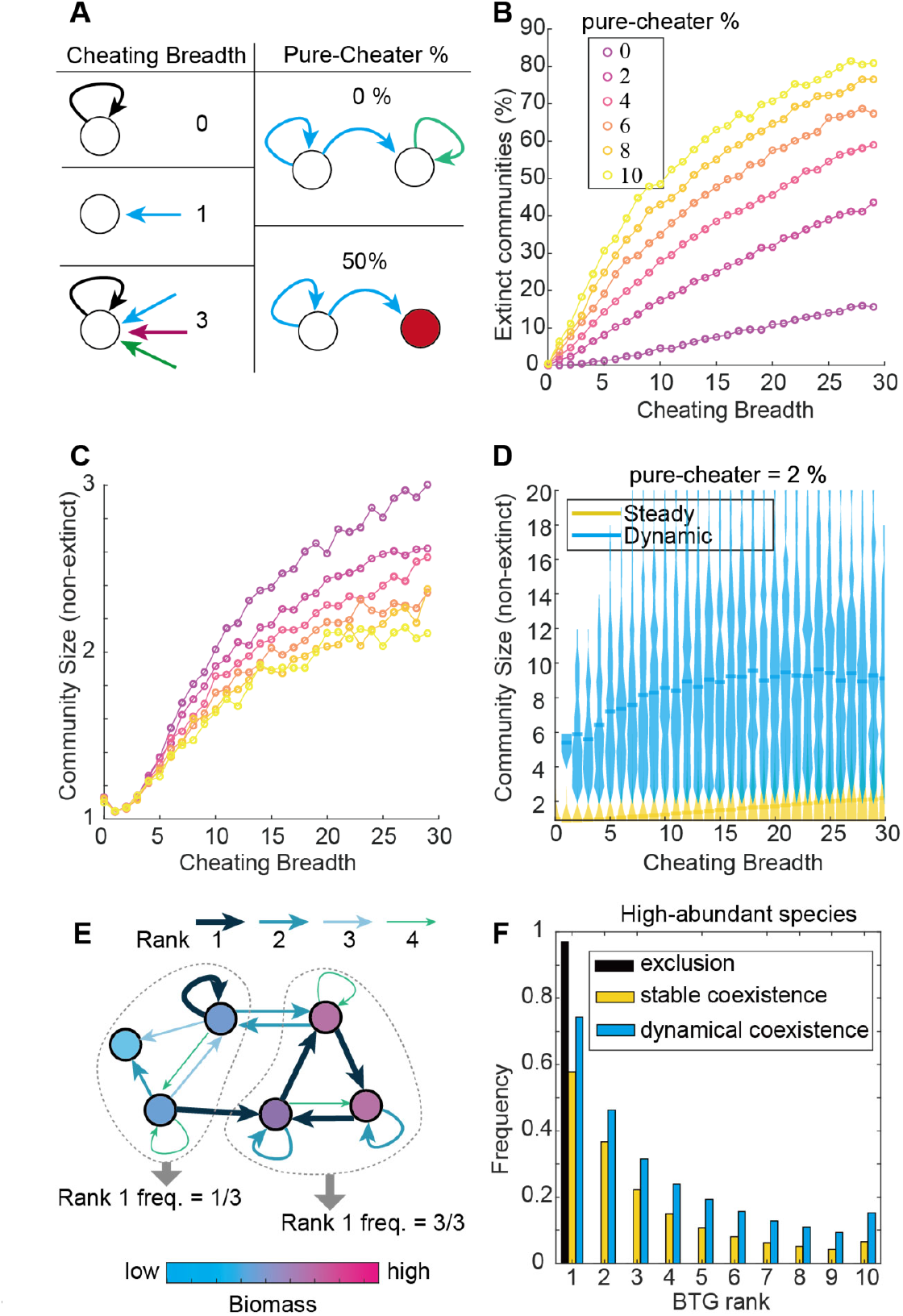
Cheating breadth elevates both extinction risk and community diversity. A. Definitions of cheating breadth (number of exploitable foreign siderophore types) and pure-cheater ratio. B. Probability of community-level extinction increases with both cheating breadth and pure-cheater ratio (ratio is color-coded; consistent across panels B–C). C. In non-extinct communities, species richness increases with cheating breadth but decreases with pure-cheater ratio. D. Violin plots showing non-extinct community size distributions for steady (yellow) versus dynamic (green) outcomes (pure-cheater ratio = 2%), under different cheating breadth. Dynamic communities consistently support higher biodiversity. E. Schematic illustrating edge ranking in BTGs, where edges from producers *i* are ranked by their weights *b*_*ij*_. Two subgraphs formed by low-biomass species (left) and high-biomass species (right) are bracketed by dashed lines, with the relative frequency of Rank-1 incoming edges shown below. F. Rank frequency distribution of benefit transfer edges in BTGs. Top-ranked edges were enriched in subgraphs formed by high-abundance species (biomass > 10^−3^)

On the other hand, within non-extinct communities (having at least one surviving species), cheating promotes biodiversity. Species richness increases monotonically with cheating breadth (Fig. 3C), and these diverse communities are more likely to exhibit dynamic behaviors (oscillations or chaos) (SI Appendix, Section 6). Notably, dynamic communities consistently support larger populations than stable ones, suggesting that temporal fluctuations create niches for species maintenance (Fig. 3D).

The paradox of cheating breadth implies that surviving communities must harbor non-random interaction structures that enable persistence. To identify these structures, we analyzed the Benefit Transfer Graph within each non-extinct community. For every siderophore producer *i*, we ranked all its outgoing benefit edges (*b*_*ij*_) by magnitude, designating the single strongest flow as the “Rank-1 edge” (Fig. 3E). Notably, we found that the subgraph formed by surviving species is overwhelmingly dominated by these Rank-1 incoming edges (Fig. 3F), far exceeding their frequency in randomly assembled subgraphs (SI Appendix, Section 6). This finding suggests that community fate is not determined by average interaction strengths, but by a specific topological backbone formed by maximal benefit flows.

### The Maximal Benefit Transfer Graph Resolves the Cheating Paradox

To decode the structural basis of the cheating paradox, we formalized the “Rank-1” backbone as a maximal Benefit Transfer Graph (mBTG). By retaining only the strongest outgoing benefit edge for each producer, the mBTG assumes the topology of a directed pseudoforest, a graph class where every node has at most one outgoing edge (Fig. 4A). A central feature of such graphs is their decomposition into Weakly Connected Components (WCCs), which are groups of nodes connected by paths regardless of direction. In directed pseudoforests, benefit flows within each WCC inevitably converge to a unique “Core”: the minimal subset of nodes with no outgoing edges to the rest of the graph (Fig. 4B).

**Figure 4.**
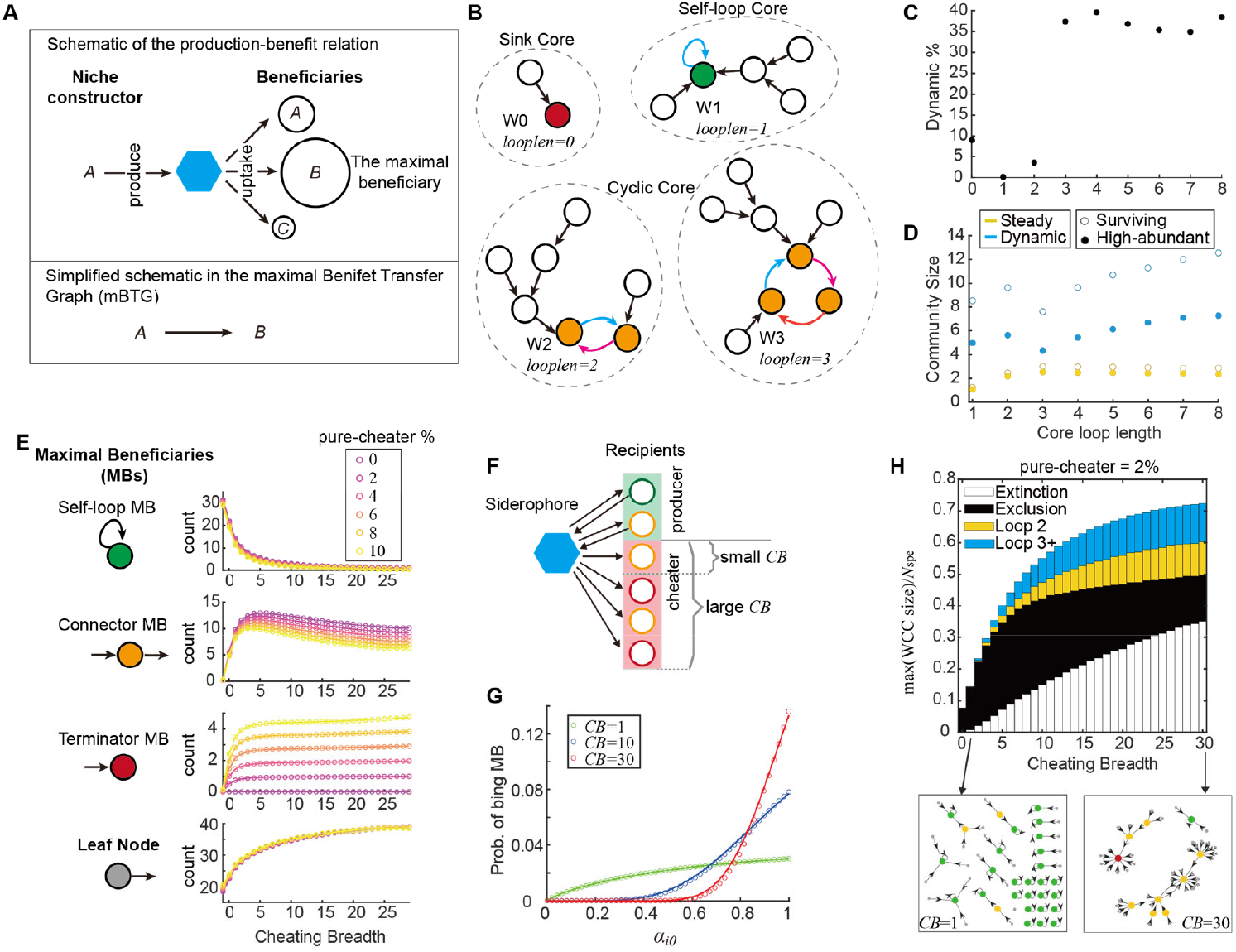
Core loops of the maximal Benefit Transfer Graph (mBTG) predicts community fate and resolves the cheating paradox. A. Construction of the mBTG. For each producer, the single strongest outgoing benefit flow defines the “maximal beneficiary,” forming a rank-1 directed edge. B. Topology of the mBTG decomposes into Weakly Connected Components (WCCs), each containing exactly one “Core” (colored nodes and edges). Four WCCs are separated by dashed circles. Core classes (Sink, Self-loop, Cyclic) are marked on each top. C. Scatter plots showing the probability of entering dynamic attractors leaps at core loop length of three. D. Scatter plot showing how community size increases with core loop length, for steady-state (yellow) and dynamic (blue) communities. Filled and open circles indicate high-abundance and surviving species (biomass threshold 10^−3^ and 10^−6^), respectively. E. Node classifications in mBTG (left) and how their counts change with cheating breadth (right). Self-loop maximal beneficiaries (MBs) have edges directed to itself; Connector MBs possess both incoming and outgoing edges; Terminator MBs have only incoming edges and no outgoing edges; Leaf nodes have no incoming edges. F. Probabilistic explanation that increasing cheating breadth expands the pool of potential recipients, diluting the producer’s chance of retaining its own siderophore (Self-loop MBs decline). G. Broader cheating amplifies MB selection bias toward high-*α*_0_ species, creating heavy-tailed in-degree distributions and promoting Terminator MBs (*CB* is abbreviation for cheating breadth). H. The percolation transition. The curve shows the fraction of nodes occupied by the largest WCC, which grows with cheating breadth. Colors under the curve indicate the proportion of WCCs governed by different core lengths. The system transitions from fragmented Self-loop Cores (exclusion) to giant components dominated by either Terminators (extinction) or Cyclic Cores (coexistence), with examples shown in bottom insets.

We discovered that the topology of this Core serves as a potent predictor of community fate, achieving arround 80% classification accuracy. Irrespective of initial conditions, asymptotic biomass consistently concentrates within a single WCC, and specifically within its core (SI Appendix, Section 7). This indicates that each WCC represents a distinct basin of attraction, with the core acting as the structural and dynamic nucleus. Characterizing these cores by their “loop length” (the number of edges forming the core) reveals three distinct architectures that dictate ecological outcomes (Fig. 4B):

1. Sink Core (Loop length 0, W0 in Fig. 4B): A core consisting of a single pure-cheater acts as a resource “black hole,” absorbing benefits without reciprocation. This topology leads to community extinction with 97% probability.
2. Self-loop Core (Loop length 1, W1 in Fig. 4B): A core formed by a producer benefiting maximally from itself leads to a steady state (Fig. 4C) where the core species survives alone, driving all others to exclusion (Fig. 4D).
3. Cyclic Core (Loop length ≥: 2, W2 and W3 in Fig.4B): A closed loop of two or more species supports coexistence. Notably, as the loop length reaches three, the probability of entering oscillatory or chaotic attractors increases sharply (Fig. 4C). While species richness in steady-state systems plateaus at short loop lengths, dynamic systems continuously support higher diversity (Fig. 4D). These dynamic communities often harbor twice as many total survivors (biomass >10^−6^) as high-abundance species (biomass > 10^−3^), suggesting that oscillatory dynamics enable rare species to persist by intermittently accessing benefits from abundant ones (Fig. 4D).

This topological framework resolves the paradox of cheating breadth. The mBTG is composed of four functional node types, whose abundance changes by cheating breadth (Fig. 4E, SI Appendix, Section 8):

1. “Self-loop” Maximal Beneficiaries (MBs): Producers that gain the highest benefit from their own siderophores. These form Self-loop Cores, driving exclusion. However, increasing cheating breadth expands the pool of potential recipients, “diluting” a producer’s probability of retaining its own siderophore as the maximal benefit (Fig. 4F). This causes a monotonic decline in exclusion-driving Self-loops.
2. “Connector” MBs: Partial-producers possessing both incoming edges and an outgoing edge. This type benefits most from foregin siderophores while producing a siderophore whose maximal beneficiary is not itself. They are the essential “glue” for Cyclic Cores, linking species into loops that support stable or dynamic coexistence.
3. “Terminator” MBs: Nodes with incoming edges but no outgoing edges, corresponding exclusively to pure-cheaters. They form Sink Cores. Selection of maximal beneficiaries probabilistically favors species with higher growth allocation (like pure-cheaters with *α*_0_ = 1), and broader cheating amplifies this bias (Fig. 4G), causing a surge in Terminator MBs.
4. “Leaf” Nodes: Species with no incoming edges. The number of Leaf nodes increases as cheating breadth expands, indicating that incoming edges concentrate disproportionately on a small subset of species. This trend can also be explained by the increased bias towards high-*α*_0_ species, which leads to a heavy-tailed in-degree distribution in which a small set of “super-beneficiaries” capture maximal benefits from many sources whereas most species get none.

Taken together, as cheating breadth increases, the mBTG undergoes a percolation-like transition. The suppression of Self-loops releases outgoing edges, while the concentration of edges toward “super-beneficiaries” promotes the coalescence of fragmented components into a giant connected cluster (Fig. 4H). This reduces multi-stability, forcing the community into few dominant attractors. Thus, while broad cheating eliminates the self-loops that trigger exclusion, it forces the community to “gamble”: it either collapses into a Terminator Sink or stabilizes into a complex, high-diversity Cyclic Core. Cheating therefore does not always act as a destabilizing vulnerability, but can also provide the necessary architectural scaffold for biodiversity.

## Discussion

In this study, we introduce an integrated framework that bridges molecular specificity, ecological dynamics, and network topology to resolve the “cheating paradox” in microbial communities. By mapping siderophore-mediated interaction dynamics to a maximal Benefit Transfer Graph (mBTG), we demonstrate that community fate, whether extinction, exclusion, or coexistence, is encoded in the core topology of benefit flows. Our findings challenge the classical view that cheating is purely detrimental. While unchecked cheating (Sink Cores) indeed drives collapse, structured exploitation (Cyclic Cores) acts as a necessary architectural scaffold for biodiversity.

The concept of niche construction traditionally emphasizes how organisms modify environments to their benefit, but it also creates vulnerabilities to exploitation[21]. While mechanisms such as spatial segregation[22], metabolic cross-feeding[23], and kin selection[24] have been proposed to resolve this dilemma, our model demonstrates that cheating itself can be transformed into an organizing force. This can be analogous to the central tenet of Modern Coexistence Theory, which posits that stable coexistence arise from either niche differentiation or fitness equivalence [25, 26]. Similarly, when microbes create multiple “chemical niches” by diverse siderophore types, partial-producers emerge as essential “loop connectors,” linking distinct niches through their dual capacity to produce and exploit. This aligns with empirical observations in *Pseudomonas*, where partial-producers are associated with diverse, non-pathogenic communities, whereas pure strategies are linked to low diversity and pathogenicity[6].

This graph-theoretic framework bridges experiments, bioinformatics, and ecological theories. While kinetic parameters are often elusive in wild communities, the structure of the Benefit Transfer Graph can be inferred from genomic analysis[4] or cross-feeding assays[27]. By coarse-graining molecular specificity into topological motifs (WCCs and Cores), the mBTG approach can forecast community fate not only in systems governed by siderophore-mediated interactions, but also in other systems driven by shared chemical resources, such as extracellular enzyme hydrolysis[28], antibiotic degradation[29], or metabolic cross-feeding networks[30].

Percolation theory describes abrupt transitions from local connectivity to global connectivity [31]. Previously applied to physical connectivity like fragmented habitats [32, 33], percolation concepts now extend to abstract interaction networks [34]. Our Maximal Benefit Transfer Graph (mBTG) exhibits distinctive percolation behavior shaped by its pseudoforest topology: increasing cheating breadth rapidly merges Weakly Connected Components into a giant component, while Strongly Connected Components (formed the cores) approach but never reach the percolation threshold. Under evolutionary dynamics, this structure self-organizes further. Pure-cheaters face extinction, driving the network toward self-sustaining structures composed by either Self-loops or Connector-based cycles. This mirrors autocatalytic set emergence, where mutually reinforcing cycles evolve spontaneously from random networks [35]. Biologically, this suggests that ecosystems operate near a critical point. Bacteria generally possess multiple siderophore receptors, ranging from 1–4 in *E. coli* to 20–30 in *Pseudomonas* and other environmental strains [7, 36-38]. This range corresponds to the “near-critical spot” in our simulations—high enough to ensure global connectivity and diversity, yet structured enough to maintain stability.

Our work provides a mechanistic foundation that grounds abstract ecological theory in molecular specificity. We show that siderophore-mediated interactions generate rich phenomena, ranging from synergistic rescue and priority effects to heteroclinic bifurcations. While phenomenological models highlight the role of intransitive interactions in maintaining diversity[39], our work identifies a concrete chemical basis for such dynamics. Furthermore, the emergence of oscillation and chaos supports theoretical predictions that non-equilibrium dynamics promote diversity through temporal niche partitioning[40].

Our current framework relies on simplified assumptions to ensure analytical tractability, such as well-mixed environments and single-siderophore production. These simplifications point toward fruitful avenues for future exploration. For instance, incorporating spatial structure could reveal how biofilm stabilize or fragment large interaction loops [41], and integrating more interactions like nutrient competition and antibiotic antagonism would offer a more holistic ecological picture [42]. Additionally, incorporating evolutionary dynamics will be crucial to understand the selective origins of these topological motifs [8, 43]. Ultimately, these future complexities will build upon the fundamental principle established here: molecular specificity transforms the social dilemma of public goods, recasting cheating from a destabilizing threat into the structural scaffold of diverse ecosystems.

## Acknowledgments

This work is supported by National Natural Science Foundation of China (No. T2321001 to ZL, 32588202 and 32425036 to SW, and T2422010 and 62172170 to XL), Fundamental and Interdisciplinary Disciplines Breakthrough Plan of the Ministry of Education of China (JYB2025XDXM502 to ZL), National Key Research and Development Programme of China (2022YFF0802103 to SW), the Peking-Tsinghua Center for Life Sciences, and the Fundamental Research Funds for Central Universities.

