## Supplementary material for "Benefit Transfer Loops Turn Cheating into a Scaffold for Microbial Diversity": SI Appendix

**This PDF file includes:**

Table S1-S2

Supporting Text Section 1-8

Figs. S1 to S15

SI Reference

**Table S1: Mathematical Symbols**

| Symbols | | Definition |
|---|---|---|
| State variables | $M_i$ | Biomass concentration of microbial species $i$ |
| | $R_j$ | Concentration of siderophore type $j$ |
| | $R_{\mathrm{iron}}$ | Free iron concentration in the medium |
| Chemostat parameters | $R_{\text{supply}}$ | Fixed iron concentration in the in-flux medium |
| | $d$ | Chemostat dilution rate |
| Microbial strategy parameters | $\alpha_{i0}$ | Fraction of resources allocated to growth by species $i$ |
| | $\alpha_{ij}$ | Fraction of resources allocated by species $i$ to produce siderophore type $j$ ($j$ >0) |
| | $v_{ij}$ | Fraction of receptors in species $i$ allocated to siderophore type $j$ |
| Community parameters | $N_{\text{spe}}$ | Total number of microbial species |
| | $N_{\text{sid}}$ | Total number of siderophore types |
| | $\sigma$ | Migration-in rate for microbes |
| | $u_j$ | Iron-binding affinity of siderophore type $j$ |
| | $\epsilon_j$ | Conversion efficiency of resources to siderophore type $j$ |
| | $\gamma$ | Efficiency of converting resources to biomass. |
| | $CB$ | Cheating breadth, defined by the average number of cheating receptors per microbe |
| | $r_c$ | Ratio of pure-cheaters in the community |

#### Table S2: Parameter Choices

<table>
<tr><td>Figure 2B, S1</td><td>$R_{\text{supply}} = 0.8, d = 0.1;\ \alpha_{10} = 0.6, v_{11} = 1;\ \sigma = 0, u = 1, \epsilon = 1, \gamma = 1$</td></tr>
<tr><td>Figure 2D, S2</td><td>$R_{\text{supply}} = 0.8, d = 0.1;\ v_{11} = v_{22} = 1;\ \sigma = 0, u = 1, \epsilon = 1, \gamma = 1$</td></tr>
<tr><td>Figure 2E-G</td><td>$R_{\text{supply}} = 0.8, d = 0.1;\ \sigma = 0, u = 1, \epsilon = 1, \gamma = 1$ , Other parameters are specified on the top of the plots</td></tr>
<tr><td>Figure 2I, S3-S4</td><td>$R_{\text{supply}} = 1, d = 0.1;\ \sigma = 0, u = 1, \epsilon = 10, \gamma = 1$ , $\alpha_{11} = 0.3, \alpha_{22} = 0.4, \alpha_{33} = 0.5;\ v_{123} = 0.7; v_{321} = 0$. Other parameters are specified on the top of the plots</td></tr>
<tr><td>Figure 2K, S5</td><td>$R_{\text{supply}} = 1, d = 0.1;\ \sigma = 0, u = 1, \epsilon = 10, \gamma = 1,\ \alpha_{ii} = 0.2$ for $i = 1 \dots 5$.<br>$v_{ij}$=
<table>
<tr><td>0.02</td><td>0.67</td><td>0</td><td>0.31</td><td>0</td></tr>
<tr><td>0</td><td>0.02</td><td>0.55</td><td>0</td><td>0.43</td></tr>
<tr><td>0.55</td><td>0</td><td>0.02</td><td>0.43</td><td>0</td></tr>
<tr><td>0</td><td>0.43</td><td>0</td><td>0.02</td><td>0.55</td></tr>
<tr><td>0.67</td><td>0</td><td>0.31</td><td>0</td><td>0.02</td></tr>
</table>
Other parameters are specified on the top of the plots</td></tr>
<tr><td>Figure 3-4, S6-11</td><td>$R_{\text{supply}} = 3, d = 0.1;\ \sigma = 10^{-8}, u = 1, \epsilon = 10, \gamma = 1$</td></tr>
</table>

#### Data Availability

All computer code and scripts used for the simulations, analysis, and figure generation in this study have been deposited in the Zenodo repository: (https://zenodo.org/records/18158439).

This study is a theoretical work and did not generate new biological materials or empirical datasets; all data supporting the findings are derived from the model simulations and are available within the article and its Supplementary Information.

### Section 1: Model Definition

Our mathematical framework for siderophore-mediated interactions comprises two interconnected components: (1) a dynamic model quantifying feedbacks between microbial populations and their chemical environment (Fig. 1A–B); and (2) a graph representation of benefit transfers between microbes via siderophore production and uptake (Fig. 1C).

#### Dynamic Model

This dynamic model extends the classical consumer-resource framework[1-3] by explicitly incorporating the active role of microbes as both consumers of iron and producers of siderophores. The model tracks three sets of variables: (1) Microbial biomass concentration $M_i$ for each species $i = 1,2,\ldots,N_{\text{spe}}$; (2) concentrations of each siderophore type $R_j$ $(j = 1,2,\ldots,N_{\text{sid}})$; (3) Free iron concentration $R_{\text{iron}}$:

$$\frac{dM_i}{dt} = M_i \cdot \left( \gamma \cdot \alpha_{i0} \cdot \sum_j v_{ij} \cdot J_j - d \right) + \sigma, \quad \text{where } i = 1,\cdots,N_{\text{spe}}. \tag{S1}$$

$$\frac{dR_j}{dt} = \sum_i M_i \cdot \alpha_{ij} \cdot \epsilon_j - d \cdot R_j, \quad \text{where } j = 1,\cdots,N_{\text{sid}}. \tag{S2}$$

$$\frac{dR_{\text{iron}}}{dt} = d \cdot \left( R_{\text{supply}} - R_{\text{iron}} \right) - \sum_{i,j} M_i \cdot v_{ij} \cdot J_j. \tag{S3}$$

This model incorporates three key assumptions:

1. **Dual resource allocation constraints**.
   1.1 Trade-off between growth and siderophore production. Each species $i$ allocates limited resources between primary metabolism for growth (fraction $\alpha_{i0}$) and production of type-$j$ siderophore (fraction $\alpha_{ij}$ for $j > 0$), satisfying:

$$\alpha_{i0} + \sum_{j=1}^{N_{\text{sid}}} \alpha_{ij} = 1. \tag{S4}$$

   1.2 Trade-off in receptor preference. Given the limited area of membrane surface and the cost of active transport, we set the fraction of receptors in species $i$ utilizing siderophore of type $j$ as $v_{ij}$, satisfying:

$$\sum_{j=1}^{N_{\text{sid}}} v_{ij} = 1. \tag{S5}$$

For each species $i$, receptors matching its own siderophores are termed “self-receptors” ( $j$ where $v_{ij} > 0$ and $\alpha_{ij} > 0$ ), while those that exploit siderophores produced by others are “cheating-receptors” ($j$ where $v_{ij} > 0$ but $\alpha_{ij} = 0$). These receptors are also known as “heterologous receptors” or “xenosiderophore receptors” from biological perspective[4]. This definition distinguishes self-reliant and exploitative iron-acquisition strategies.

2. **Iron as the sole limiting resource**. For simplification, we focus exclusively on iron competition. Facilitation of iron uptake by the siderophore-iron complex of type $j$, $J_j$, is approximated by mass-action kinetics, assuming rapid equilibrium in complex formation:

$$J_j = u_j \cdot R_j \cdot R_{\text{iron}}, \tag{S6}$$

where $u_j$ denotes the affinity of siderophore $j$ for iron.

Each species’ growth rate scales linearly with the total iron uptake ($\sum_{j=1}^{N_{\text{sid}}} v_{ij} J_j$) and its growth allocation fraction ($\alpha_{i0}$):

$$growth_i = \gamma \cdot \alpha_{i0} \cdot \sum_{j=1}^{N_{\text{sid}}} v_{ij} \cdot J_j, \tag{S7}$$

with $\gamma$ representing the efficiency of resource-to-growth conversion.

(3) **Chemostat Dynamics**: The environment is modeled as a chemostat with dilution rate $d$ and iron supply concentration $R_{\text{supply}}$. Microbes grow according to Eq.(S7), are diluted at rate $d$, with a small immigration rate $\sigma$ (Eq.(S1)). Each microbial species $i$ produces siderophore type $j$ at the rate proportional to $\alpha_{ij}$, with a conversion factor $\epsilon_j$ (Eq. (S2)). Iron, the sole external resource, is supplied at concentration $R_{\text{supply}}$, and consumed by microbes with the assistance of siderophores (Eq. (S3)).

### Strategy Space of Microbial Iron Acquisition

The resource allocation parameters define three classes of siderophore strategies (Fig. 1B):

(1) Pure-producers (Fig. 1B, left panel): synthesize and exclusively utilize their own siderophore $j$, ($\alpha_{ij} > 0$ and $v_{ij} = 1$); classical “cooperators” in game theory [5].

(2) Partial-producers (Fig. 1B, middle panel): both produce their own siderophores and maintain the capacity to uptake siderophores produced by other species ($\alpha_{ij} > 0, v_{ij} < 1$ for some $j$, and there exists a $k$ such that $\alpha_{ik} = 0, v_{ik} > 0$). This strategy, commonly observed among environmental *Pseudomonas* strains, represents a balance between self-sufficiency and opportunistic resource exploitation[6].

(3) “Pure-cheaters” (Fig. 1B, right panel): invest fully in growth ($\alpha_{i0} = 1$) and exploit siderophores produced by others ($v_{ij} > 0$ for some $j$); classical “cheater” in game theory [7].

#### Definition of the Benefit Transfer Graph

To quantify the interactions mediated by siderophores, we also defined the "Benefit Transfer Graph (BTG)" describing how siderophore production by one species benefits others (Fig. 1C). When microbe $m1$ produces siderophores of type $j$, all species with matching receptors ($v_{ij} > 0$) can benefit. These benefits are proportional to the shared environmental siderophore-iron complex represented by $J_j$ (Eq.(S6)), as well as species-specific growth allocation $\alpha_{i0}$ and receptor allocation $v_{ij}$. Thus, the benefit transferred from producer $m1$ to recipient $m2$, per biomass for both species, is:

$$b_{m1,m2} = \alpha_{m2,0} \sum_{j=1}^{N_{\mathrm{sid}}} \left( v_{m2,j} \cdot \alpha_{m1,j} \right) \tag{S8}$$

In the limiting case that each species produces no more than one type of siderophore, this simplifies to:

$$b_{m1,m2} = \begin{cases} 0, & \text{if } \alpha_{m1,j} = 0 \text{ for all } j \\ \alpha_{m2,0} \cdot v_{m2,j}, & \text{if } \alpha_{m1,j} > 0 \end{cases}$$

All pairwise benefit coefficients $b_{m_1,m_2}$ together define a Benefit Transfer Graph (BTG), a directed graph in which nodes represent microbial species and edges represent benefit flows from siderophore producers to recipients. An edge from node $m_1$ to node $m_2$ exists when species $m_2$ gains growth benefits from siderophores produced by $m_1$ (i.e., $b_{m_1,m_2} > 0$).

Under the reasonable assumption that each species can utilize its own siderophore ($v_{ij} > 0$ whenever $\alpha_{ij} > 0$), every producer species has a self-loop in the BTG (Fig. 1C, first and second panels). When a species exploits siderophores produced by another, a directed edge points from the producer to the cheater (Fig. 1C, third and last panels). This Benefit Transfer Graph provides an intuitive depiction of community iron interactions: who produces the siderophores and who benefits from them.

### Section 2: One Pure-Producer Exhibits Bistability

In a single-species system (Fig. 2A), siderophore secretion generates a positive feedback loop: siderophores enhance iron uptake, which promotes growth and further siderophore production. In this system, Eq.(S1)-(S3) reduce to a three-variable system:

$$\frac{dM_1}{dt} = M_1 \cdot (\gamma \cdot \alpha_{10} \cdot v_{11} \cdot u_1 \cdot R_1 \cdot R_{\text{iron}} - d), \tag{S9}$$

$$\frac{dR_1}{dt} = M_1 \cdot (1 - \alpha_{10}) \cdot \epsilon_1 - d \cdot R_1, \tag{S10}$$

$$\frac{dR_{\text{iron}}}{dt} = d \cdot \left(R_{\text{supply}} - R_{\text{iron}}\right) - M_1 \cdot v_{11} \cdot u_1 \cdot R_1 \cdot R_{\text{iron}}. \tag{S11}$$

Microbial growth depends on the concentrations of siderophore ($R_1$) and iron ($R_{\text{iron}}$), thus these defining a two-dimensional chemical space $[R_1, R_{\text{iron}}]$ that directly interacting with the species. Each point in this space represents a specific chemical environment.

#### Steady-State Analysis

We conducted a steady-state analysis, where steady-state values are indicated by "*". At steady-state, $\frac{dM_1^*}{dt} = 0$ in Eq. (S9) describes the growth contour, representing all chemical environments where the microbial growth rate equals the dilution rate [8]:

$$\gamma \cdot \alpha_{10} \cdot v_{11} \cdot u_1 \cdot R_1^* \cdot R_{\text{iron}}^* = d. \tag{S12}$$

This equation describes a hyperbolic relationship in the chemical space (**Figure S1**A, black curve):

$$R_{\text{iron}}^* = \frac{d}{\gamma \cdot v_{11} \cdot u_1 \cdot \alpha_{10}} \cdot \frac{1}{R_1^*}, \tag{S13}$$

At the steady state, $\frac{dR_1^*}{dt} = 0$ in Eq. (S10) leads to a linear relationship between $M_1^*$ and $R_1^*$:

$$M_1^* = \frac{d}{(1 - \alpha_{10}) \cdot \epsilon_1} \cdot R_1^*. \tag{S14}$$

Similar, for the steady-state of iron, setting $\frac{dR_{\text{iron}}^*}{dt} = 0$ in Eq. (S11) gives:

$$M_1^* = d \cdot \frac{R_{\text{supply}} - R_{\text{iron}}^*}{u_1 \cdot v_{11} \cdot R_1^* \cdot R_{\text{iron}}^*}. \tag{S15}$$

Substituting the relationship $\gamma \cdot \alpha_{10} \cdot u_1 \cdot v_{11} \cdot R_1^* \cdot R_{\text{iron}}^* = d$ from Eq. (S12) into Eq. (S15), we obtain another linear relationship between $M_1^*$ and $R_{\text{iron}}^*$:

$$M_1^* = \left(R_{\text{supply}} - R_{\text{iron}}^*\right) \cdot \gamma \cdot \alpha_{10}. \tag{S16}$$

By combining Eq. (S14) and (S16), the flux balance requirement in steady states is

$$\frac{d}{(1-\alpha_{10})\cdot\epsilon_1}\cdot R_1^* = \left(R_{\text{supply}} - R_{\text{iron}}^*\right)\cdot\gamma\cdot\alpha_{10}. \tag{S17}$$

This leads to a decreasing linear relationship between $R_{\text{iron}}^*$ and $R_1^*$, where higher $R_1^*$ corresponds to higher $M_1^*$ (colored dot line in **Figure S1**A):

$$R_{\text{iron}}^* = R_{\text{supply}} - \frac{d}{\gamma\cdot\alpha_{10}\cdot(1-\alpha_{10})\cdot\epsilon_1}\cdot R_1^* \tag{S18}$$

The intersections of the growth contour (Eq. (S13)) and the flux-balance line (Eq. (S18)) represent the system's fixed points, which are necessary for steady-state conditions (**Figure S1**A). Graphically, the number of intersections depends on the relative position of the growth contour and the flux-balance line. For example, increasing $R_{\text{supply}}$ shifts the y-axis intercept of the flux-balance line upward, increasing the likelihood of two intersections.

Additionally, the system always has a "full-extinction" state, where no microbes grow, and no siderophores are produced. This state satisfies the fixed-point conditions derived from Eq. (S9)-(S11):

$$M_1 = 0, R_1 = 0, R_{\text{iron}} = R_{\text{supply}} \tag{S19}$$

The extinction state corresponds to the leftmost point on the flux-balance line and represents a scenario where microbial growth cannot be sustained. It is always stable for any systems satisfying Eq. (1)-(3)

#### Analytical Solutions of Fixed Points

Solving Eq. (S13) and (S18) results in a univariate quadratic equation for $R_1^*$:

$$\frac{d}{\gamma\cdot\alpha_{10}\cdot(1-\alpha_{10})\cdot\epsilon_1}{R_1^*}^2 - R_1^*\cdot R_{\text{supply}}^* + \frac{d}{\gamma\cdot v_{11}\cdot u_1\cdot\alpha_{10}} = 0 \tag{S20}$$

This quadratic equation has two solutions for $R_1^*$, provided the discriminant is non-negative:

$$R_1^* = \frac{R_{\text{supply}} \pm \sqrt{{R_{\text{supply}}}^2 - 4\cdot\frac{d^2}{\gamma^2\cdot\alpha_{10}^2\cdot(1-\alpha_{10})\cdot\epsilon_1\cdot v_{11}\cdot u_1}}}{2\cdot\frac{d}{\gamma\cdot\alpha_{10}\cdot(1-\alpha_{10})\cdot\epsilon_1}}. \tag{S21}$$

These solutions are meaningful (i.e., real and positive) only under the condition:

$$\gamma\frac{R_{\text{supply}}}{2d} > \frac{1}{\alpha_{10}\sqrt{(1-\alpha_{10})\cdot\epsilon_1\cdot v_{11}\cdot u_1}} \tag{S22}$$

In summary, the dynamic system described by Eq. (S9)-(S11) can have either one or three fixed point, depending on whether the condition in Eq. (S22) is satisfied.

#### Bifurcation Analysis

When $R_{\text{supply}}$ is too small or $d$ is too large, the condition in Eq. (S22) is not satisfied. In this case, the growth contour and the flux-balance line do not intersect, resulting in a system that only has the full-extinction state (Eq. (S19)).

However, when $R_{\text{supply}}$ is sufficiently large or $d$ is small enough to satisfy Eq. (S22), the growth contour and the flux-balance line intersect, yielding three fixed points. In this scenario, the extinction state remains stable, and the fixed point with the higher $R_1^*$ value is also stable, while the intermediate intersection is unstable. Therefore, the single-species system exhibits bi-stability: higher initial biomass or siderophore concentration allows the microbe to thrive, whereas insufficient initial biomass (below a critical threshold) leads to extinction, analogous to the Allee effect in ecology [9].
Changes in environmental factors, such as variations in iron supply concentration $R_{\text{supply}}$ and dilution rate $d,$ can trigger bifurcations in the system. Intuitively, when the $R_{\text{supply}}$ is low, the system only has the extinction state. As $R_{\text{supply}}$ increases, the system undergoes a saddle-node bifurcation, transitioning to a bistable regime. In this regime, microbes with sufficiently high initial biomass can successfully establish themselves in the niche (**Figure S1**B).

It is also important to note that the resource allocation term $\alpha_{10}$ also influences whether the condition in Eq. (S22) is satisfied. If $\alpha_{10}$ is either too small or too large (indicating that cells allocate too much or too little to siderophore production), the denominator in Eq. (S22), becomes small, pushing the system beyond the threshold and into the total extinction regime (**Figure S1**C).

### Section 3: Two Pure-Producers Cannot Coexist

We begin by considering the simplest two-species system (**Figure S2**A), which consists of two pure-producers (species 1 and 2). Each species produces a distinct type of siderophore (type 1 and type 2, respectively) and exclusively uptakes its own siderophore, without cheating siderophores from each other ($v_{11} = 1,\ v_{22} = 1$).
For species 2, similar to the dynamics of species 1 described in Eq. (S9)-(S10), the biomass and siderophore dynamics are governed by the following equations:

$$\frac{dM_2}{dt} = M_2 \cdot (\gamma \cdot \alpha_{20} \cdot v_{22} \cdot u_2 \cdot R_2 \cdot R_{\text{iron}} - d), \tag{S23}$$

$$\frac{dR_2}{dt} = M_2 \cdot (1 - \alpha_{20}) \cdot \epsilon_1 - d \cdot R_2. \tag{S24}$$

Since both species can uptake iron, Equation (S11) is modified to account for the iron uptaking from two species, resulting in the following updated equation for free iron:

$$\frac{dR_{\text{iron}}}{dt} = d \cdot \left(R_{\text{supply}} - R_{\text{iron}}\right) - M_1 \cdot u_1 \cdot v_{11} \cdot R_1 \cdot R_{\text{iron}} - M_2 \cdot u_2 \cdot v_{22} \cdot R_2 \cdot R_{\text{iron}}. \tag{S25}$$

In this two-species system described by Eq. (S9)-(S10) and Eq. (S23)-(S25), the species interact solely through competition for iron, as each species depletes the available iron via its siderophore production.

#### Phase Diagram Analysis

Phase diagram of this system shows that coexistence is impossible across all parameters (**Figure S2**B). First, due to the Allee effect described above, a steady-state of “all species extinct” remains stable for this and for all other systems in this work. Beyond this trivial attractor, each species’ survival solely depends on its own growth allocation (**Figure S2**B), dividing the $\alpha_{10}$-$\alpha_{20}$ parameter space into four kinds of regimes. In the global extinction regime (0) (Fig. 2G, first panel), neither species satisfies the survival condition in Eq.(S22), and the full extinction steady-state is the only attractor. At the two dominance regions (1) and (2), one species possesses a distinct fitness advantage and deterministically excludes the other regardless of initial abundance (Fig. 2G, second panel). The regime of priority effect (3) appears when both species satisfy the bistability requirement, with three attractors in the state-space: species m1 dominates, species m2 dominates, or both go extinct. The final state depends on which species is initially more dominant (Fig. 2G, third panel).

#### Graphical Representation of Competitive Exclusion

Using graphical representation in chemical space, we can intuitively understand why these two species cannot coexist. As shown in **Figure S2**C, when species 1 and species 2 are grown alone, they create distinct chemical environments with different free iron concentrations. For example, a small population of species 1 cannot invade the environment established by species 2, because there is no siderophore type 1 available for species 1 to utilize. This results in zero growth rate for species 1, as predicted by Equation (S9). Similarly, species 2 cannot invade the environment created by species 1, as that environment lacks siderophore type 2.

Now, consider a hypothetical scenario where the two species might coexist at equilibrium ($M_1 > 0$ and $M_2 > 0$, with Equation (S9)-(S10) and (S23)-(S25) are all set to zero). From the perspective of species 1, at steady state, the effective iron supply is reduced by species 2's siderophore production, from $R_{\text{supply}}$ into $R_{\text{supply}} - \frac{M_2}{\gamma\alpha_{20}}$. In this case, a small increase in $M_2$ further reduces the effective iron supply for species 1, thereby decreasing its growth rate and, consequently, its biomass $M_1$. Conversely, for species 2, the effective iron supply concentration is $R_{\text{supply}} - \frac{M_1}{\gamma\alpha_{10}}$, A reduction in $M_1$ leads to an increase in the growth rate and biomass of species 2. This feedback loop amplifies small perturbations in $M_2$, making the equilibrium unstable.

Thus, despite the system having three "dimensions" in the chemical space, [$R_{\text{supply}}$, $R_1$, $R_2$], the species still exhibit competitive exclusion, meaning that no more than one species can stably survive in the system.

### Section 4: Conditions for Stable Coexistence Between Two Partial-Producers

Next, we derived the condition for coexistence of two partial-producers (species 1 and 2), each producing a distinct type of siderophore (concentrations denoted as $[R_1]$ and $[R_2]$, respectively). In this system (Fig. 2C), each species uptakes its own siderophore as well as the siderophore produced by the other species (i.e., $v_{12} > 0,\ v_{21} > 0$). The dynamics of the five variables in this system are described by the following equations:

$$\frac{dM_1}{dt} = M_1 \cdot (\gamma \cdot \alpha_{10} \cdot (u_1 \cdot v_{11} \cdot R_1 + u_2 \cdot v_{12} \cdot R_2) \cdot R_{\text{iron}} - d). \tag{S26}$$

$$\frac{dR_1}{dt} = M_1 \cdot (1 - \alpha_{10}) \cdot \epsilon_1 - d \cdot R_1. \tag{S27}$$

$$\frac{dM_2}{dt} = M_2 \cdot (\gamma \cdot \alpha_{20} \cdot (u_1 \cdot v_{21} \cdot R_1 + u_2 \cdot v_{22} \cdot R_2) \cdot R_{\text{iron}} - d). \tag{S28}$$

$$\frac{dR_2}{dt} = M_2 \cdot (1 - \alpha_{20}) \cdot \epsilon_1 - d \cdot R_2. \tag{S29}$$

$$\frac{dR_{\text{iron}}}{dt} = d \cdot \left(R_{\text{supply}} - R_{\text{iron}}\right) - M_1 \cdot (u_1 \cdot v_{11} \cdot R_1 + u_2 \cdot v_{12} \cdot R_2) \cdot R_{\text{iron}} - M_2 \cdot (u_1 \cdot v_{21} \cdot R_1 + u_2 \cdot v_{22} \cdot R_2) \cdot R_{\text{iron}}. \tag{S30}$$

#### Analytical Solution of Four Types of Fixed-Points

At steady state (denoted by *), all rates of change in Eq. (S26)-(S30) equal zero.

From Eq. (S27) and Eq. (S29), we obtain:

$$M_1^* = \frac{d}{(1 - \alpha_{10}) \cdot \epsilon_1} \cdot R_1^*, \tag{S31}$$

$$M_2^* = \frac{d}{(1 - \alpha_{20}) \cdot \epsilon_1} \cdot R_2^*. \tag{S32}$$

For steady-state conditions ($dM_i^*/dt = 0$) from Eq. (S26) and Eq. (S28) yield four possible fixed-point scenarios:

(1) Full-Extinction: $M_1^* = 0,\ M_2^* = 0$.

(2) Species 1 exclude species 2: $M_1^* > 0$, $M_2^* = 0$. This case reduces to the single-producer scenario from Section 2 (Eq. (S13), (S14), and (S21)). The fixed-point solves to:

$$R_1^* = \frac{R_{\text{supply}} \pm \sqrt{{R_{\text{supply}}}^2 - 4 \cdot \frac{d^2}{v_{11} \cdot \gamma^2 \cdot \alpha_{10}^2 \cdot (1 - \alpha_{10}) \cdot \epsilon_1 \cdot u_1}}}{2 \cdot \frac{d}{\alpha_{10} \cdot (1 - \alpha_{10}) \cdot \epsilon_1}}, \tag{S33}$$

$$M_1^* = \frac{d}{(1-\alpha_{10})\cdot\epsilon_1}\cdot R_1^*,$$
$$R_{\text{iron}}^* = \frac{d}{\gamma\cdot u_1\cdot\alpha_{10}}\cdot\frac{1}{R_1^*},$$
$$R_2^* = 0, M_2^* = 0.$$

(3) Species 2 exclude species 1: $M_1^* = 0$, $M_2^* > 0$. Similarly to case 2, the fixed-point solves to:

$$R_2^* = \frac{R_{\text{supply}} \pm \sqrt{{R_{\text{supply}}}^2 - 4\cdot\frac{d^2}{v_{22}\cdot\gamma^2\cdot\alpha_{20}^2\cdot(1-\alpha_{20})\cdot\epsilon_1\cdot u_1}}}{2\cdot\frac{d}{\alpha_{2,0}\cdot(1-\alpha_{20})\cdot\epsilon_1}}, \quad \text{(S34)}$$
$$M_2^* = \frac{d}{(1-\alpha_{20})\cdot\epsilon_1}\cdot R_2^*,$$
$$R_{\text{iron}}^* = \frac{d}{\gamma\cdot u_1\cdot\alpha_{20}}\cdot\frac{1}{R_2^*},$$
$$R_1^* = 0, M_1^* = 0.$$

(4) Coexistence: $M_1^* > 0$, $M_2^* > 0$, which give:

$$\gamma\cdot\alpha_{10}\cdot(u_1\cdot v_{11}\cdot R_1^* + u_2\cdot v_{12}\cdot R_2^*)\cdot R_{\text{iron}}^* = d, \quad \text{(S35)}$$
$$\gamma\cdot\alpha_{20}\cdot(u_1\cdot v_{21}\cdot R_1^* + u_2\cdot v_{22}\cdot R_2^*)\cdot R_{\text{iron}}^* = d.$$

Solving for $R_1^*$ and $R_2^*$ in terms of $R_{\text{iron}}^*$, we obtain:

$$R_1^* = \frac{d}{R_{\text{iron}}^*\cdot\gamma\cdot u_1}\frac{\alpha_{10}\cdot v_{12} - \alpha_{20}\cdot v_{22}}{\alpha_{10}\cdot\alpha_{2,0}\cdot(v_{12}\cdot v_{21} - v_{11}\cdot v_{22})}, \quad \text{(S36)}$$
$${R_2}^* = \frac{d}{R_{\text{iron}}^*\cdot\gamma\cdot u_2}\frac{\alpha_{20}\cdot v_{21} - \alpha_{10}\cdot v_{11}}{\alpha_{10}\cdot\alpha_{20}\cdot(v_{12}\cdot v_{21} - v_{11}\cdot v_{22})}.$$

For the free iron concentration at fixed point, $\frac{dR_{\text{iron}}^*}{dt} = 0$ in Eq.(S30), combining Eq.(S35) gives:

$$d\cdot\left(R_{\text{supply}} - R_{\text{iron}}^*\right) = \frac{M_1^*\cdot d}{\gamma\cdot\alpha_{10}} + \frac{M_2^*\cdot d}{\gamma\cdot\alpha_{20}}. \quad \text{(S37)}$$

Substituting the biomass terms from Eq. (S31) and Eq. (S32) into Eq. (S37), we obtain the fixed-point flux balance condition with three variables $R_{\text{iron}}^*$, $R_1^*$ and $R_2^*$:

$$R_{\text{supply}} - R_{\text{iron}}^* = \frac{\frac{d}{(1-\alpha_{10})\cdot\epsilon_1}R_1^*}{\gamma\cdot\alpha_{10}} + \frac{\frac{d}{(1-\alpha_{20})\cdot\epsilon_1}R_2^*}{\gamma\cdot\alpha_{20}}. \quad \text{(S38)}$$

Substituting $R_1^*$ and $R_2^*$ from Eq. (S36) into Eq. (S38), we get the fixed-point requirement with only $R_{\text{iron}}^*$ as the variable:

$$R_{\text{supply}} - R_{\text{iron}}^{*} = \frac{1}{R_{\text{iron}}^{*}} d^2 \frac{\frac{1}{\epsilon_1 \cdot u_1}\frac{\alpha_{10}\cdot v_{12} - \alpha_{20}\cdot v_{22}}{\alpha_{10}\cdot\left(1-\alpha_{1,0}\right)} + \frac{1}{\epsilon_1 \cdot u_2}\frac{\alpha_{20}\cdot v_{21} - \alpha_{10}\cdot v_{11}}{\alpha_{20}\cdot\left(1-\alpha_{2,0}\right)}}{\gamma^2\cdot\alpha_{10}\cdot\alpha_{20}\cdot(v_{12}\cdot v_{21} - v_{11}\cdot v_{22})}. \tag{S39}$$

Introducing the hyperparameter $A$ to simplify the equation,

$$A = d^2 \frac{\frac{1}{\epsilon_1 \cdot u_1}\frac{\alpha_{10}\cdot v_{12} - \alpha_{20}\cdot v_{22}}{\alpha_{10}\cdot(1-\alpha_{10})} + \frac{1}{\epsilon_1 \cdot u_2}\frac{\alpha_{20}\cdot v_{21} - \alpha_{10}\cdot v_{11}}{\alpha_{20}\cdot(1-\alpha_{20})}}{\gamma^2\cdot\alpha_{10}\cdot\alpha_{20}\cdot(v_{12}\cdot v_{21} - v_{11}\cdot v_{22})} \tag{S40}$$

The equation then simplifies to:

$${R_{\text{iron}}^{*}}^{2} - R_{\text{supply}}\cdot R_{\text{iron}}^{*} + A = 0 \tag{S41}$$

And the fixed-point concentration of free iron, $R_{\text{iron}}^{*}$, solves to:

$$R_{\text{iron}}^{*} = \frac{R_{\text{supply}}}{2} \pm \frac{\left(R_{\text{supply}}^{2} - 4\cdot A\right)^{\frac{1}{2}}}{2} \tag{S42}$$

From the solution of $R_{\text{iron}}^{*}$ at Eq. (S42), Eq. (S36) gives out the fixed-point solution of $R_1^*$ and $R_2^*$, and Eq. (S31)-(S32) gives the steady state value of $M_1^*$ and $M_2^*$.

Taken together, for biologically meaningful coexistence fixed-point values, there are several requirements:

1. For positive $R_1^*$ and ${R_2}^*$, Eq. (S36) requires that all three following terms

$$\begin{gathered} \alpha_{10}\cdot v_{12} - \alpha_{20}\cdot v_{22}, \\ \alpha_{20}\cdot v_{21} - \alpha_{10}\cdot v_{11}, \\ v_{12}\cdot v_{21} - v_{11}\cdot v_{22} = 1 - v_{11} - v_{22} \end{gathered} \tag{S43}$$

have the same sign.

2. For a real solution in Eq. (S43) to exist, the discriminant of this quadratic equation must be positive, which requires:

$$\gamma\frac{R_{\text{supply}}}{2d} > \sqrt{\frac{\frac{1}{\epsilon_1 \cdot u_1}\frac{\alpha_{10}\cdot v_{12} - \alpha_{20}\cdot v_{22}}{\alpha_{10}\cdot(1-\alpha_{10})} + \frac{1}{\epsilon_1 \cdot u_2}\frac{\alpha_{20}\cdot v_{21} - \alpha_{10}\cdot v_{11}}{\alpha_{20}\cdot\left(1-\alpha_{2,0}\right)}}{\alpha_{10}\cdot\alpha_{20}\cdot(v_{12}\cdot v_{21} - v_{11}\cdot v_{22})}} \tag{S44}$$

### Stability Analysis of the Fixed Points

To analyze the stability of the coexistence fixed point, we first introduce the following abbreviations:

$$\begin{gathered} J_1^* = \frac{d\cdot(\alpha_{20}\cdot v_{22} - \alpha_{10}\cdot v_{12})}{\gamma\cdot\alpha_{10}\cdot\alpha_{20}\cdot(v_{11}+v_{22}-1)}, \\ J_2^* = \frac{d\cdot(\alpha_{10}\cdot v_{11} - \alpha_{20}\cdot v_{21})}{\gamma\cdot\alpha_{10}\cdot\alpha_{20}(v_{11}+v_{22}-1)}, \end{gathered} \tag{S45}$$

$$c_{i0} = M_i^* \cdot \sum_j v_{ij} \cdot \left(\frac{\partial J_j}{\partial R_{\text{iron}}}\right)^*$$

$$c_{ij} = M_i^* \cdot v_{ij} \cdot (\frac{\partial J_j}{\partial R_j})^* \text{ , (for } j = 1,2)$$

At the coexistence fixed point, we derive the Jacobian matrix:

$$\text{Jacobian} = \begin{pmatrix} 0 & 0 & \gamma \cdot \alpha_{10} \cdot c_{11} & \gamma \cdot \alpha_{10} \cdot c_{12} & \gamma \cdot \alpha_{10} \cdot c_{10} \\ 0 & 0 & \gamma \cdot \alpha_{20} \cdot c_{21} & \gamma \cdot \alpha_{20} \cdot c_{22} & \gamma \cdot \alpha_{20} \cdot c_{20} \\ (1-\alpha_{10}) \cdot \epsilon_1 & 0 & -d & 0 & 0 \\ 0 & (1-\alpha_{20}) \cdot \epsilon_2 & 0 & -d & 0 \\ -\sum_j v_{1j} \cdot J_j^* & -\sum_j v_{2j} \cdot J_j^* & -\sum_i c_{i1} & -\sum_i c_{i2} & -d - \sum_i c_{i0} \end{pmatrix}. \quad \text{(S46)}$$

For stability analysis, we examine the characteristic equation $a_0 \lambda^5 + a_1 \lambda^4 + \cdots + a_5 = 0$. According to the Routh-Hurwitz criteria, stability requires all coefficients to be positive. The key coefficients $a_5$ and $a_4$ are:

$$a_5 = d \cdot M_1^* \cdot M_2^* \cdot \gamma^2 \cdot \alpha_{10} \cdot (1-\alpha_{10}) \cdot \epsilon_1 \cdot u_1 \cdot \alpha_{20} \cdot (1-\alpha_{20}) \cdot \epsilon_2 \cdot u_2 \cdot (1 - v_{11} - v_{22}) \cdot R_{\text{iron}}^* \cdot \left(R_{\text{supply}} - 2R_{\text{iron}}^*\right),$$

$$a_4 = \left(\frac{d^4}{R_{\text{iron}}^*} + 2R_{\text{iron}}^* \frac{R_{\text{supply}} - R_{\text{iron}}^*}{d\left(R_{\text{supply}} - 2R_{\text{iron}}^*\right)^2} a_5\right) \cdot \left(R_{\text{supply}} - 2R_{\text{iron}}^*\right). \quad \text{(S47)}$$

From these expressions, we can derive necessary conditions for stability:

1. $a_5 > 0$ requires $1 - v_{11} - v_{22}$ and $R_{\text{supply}} - 2R_{\text{iron}}^*$ have the same sign.
2. $a_4 > 0$ requires that $R_{\text{supply}} - 2R_{\text{iron}}^*$ should be positive, given $R_{\text{supply}} - R_{\text{iron}}^*$ must be positive.

These conditions, combined with earlier constraints, yield the following requirements for stable coexistence:

$$R_{\text{supply}} > 2R_{\text{iron}}^*, \quad \text{(S48)}$$

and

$$1 > v_{11} + v_{22}, \quad \text{(S49)}$$
$$\alpha_{10} \cdot v_{12} > \alpha_{20} \cdot v_{22},$$
$$\alpha_{20} \cdot v_{21} > \alpha_{10} \cdot v_{11}.$$

as the necessary conditions for two partial-producers to stably coexist.

According to the requirement in Eq. (S48), the two possible values in in Eq. (S42) consolidate to :

$$R_{\text{iron}}^* = \frac{R_{\text{supply}}}{2} - \frac{\left(R_{\text{supply}}^2 - 4 * A\right)^{\frac{1}{2}}}{2}. \quad \text{(S50)}$$

Eq. (S49) fully overlaps with the non-negative requirement in Eq. (S36).

### Parameter Requirements for Coexistence, Dominance, Priority Effects, and Extinction

Taken together, for the coexistence fixed point described by Eq. (S50), (S31)-(S32) and (S36) to exist and be stable, it requires that conditions in Eq. (S44) and Eq. (S49) to be satisfied:

$$b_{2,1} > b_{2,2}, b_{1,2} > b_{1,1} \text{ (stability condition), and}$$

$$\gamma \frac{R_{\text{supply}}}{2d} > \sqrt{\frac{\frac{1}{\epsilon_1 \cdot u_1} \frac{\alpha_{10} \cdot v_{12} - \alpha_{20} \cdot v_{22}}{\alpha_{10} \cdot (1-\alpha_{10})} + \frac{1}{\epsilon_1 \cdot u_2} \frac{\alpha_{20} \cdot v_{21} - \alpha_{10} \cdot v_{11}}{\alpha_{20} \cdot (1-\alpha_{2,0})}}{\alpha_{10} \cdot \alpha_{20} \cdot (v_{12} \cdot v_{21} - v_{11} \cdot v_{22})}} \text{ (existence condition)} \quad \text{(S51)}$$

Similarly, for two single-species fixed points, we find:

1. Species 1 excluding species 2 ($M_1^* > 0, M_2^* = 0$, details in Eq. (S33)) requires:

$$b_{1,2} < b_{1,1} \text{ (stability condition), and}$$

$$\gamma \frac{R_{\text{supply}}}{2d} > \frac{1}{\alpha_{10} \sqrt{v_{11} \cdot (1-\alpha_{10}) \cdot \epsilon_1 \cdot u_1}} \text{ (existence condition)} \quad \text{(S52)}$$

2. Vice versa, Species 2 excluding species 1 ($M_2^* > 0, M_1^* = 0$, details in Eq. (S34)) requires:

$$b_{2,1} < b_{2,2} \text{ (stability condition), and}$$

$$\gamma \frac{R_{\text{supply}}}{2d} > \frac{1}{\alpha_{20} \sqrt{v_{22} \cdot (1-\alpha_{20}) \cdot \epsilon_1 \cdot u_1}} \text{ (existence condition)} \quad \text{(S53)}$$

Besides, the full-extinction state is always stable ($M_1^* = M_2^* = 0$).

Taken together, Eq.(S51)-(S53) gives out the phase-diagram of the system (Fig. 2E-G), where the five types of possible state-space are (always have the trivial full-extinction state):

(0) Global extinction: if none of the conditions in Eq.(S51)-(S53) can be satisfied, no species survive alone or together, leaving only have this extinction state.

(1) Species 1 dominance: only Eq. (S52) is satisfied. Species m1 possesses a distinct fitness advantage and deterministically excludes species m2, regardless of their initial abundance unless they extinct together.

(2) Species 2 dominance: only Eq. (S53) is satisfied. Similar to (1), but species m2 wins.

(3) Priority effect: When both Eq. (S52) and Eq. (S53) are satisfied, there are three attractors in the state-space: species m1 excluding species m2, species m2 excluding species m1, or both go extinct. The final state depends on which species is initially more dominant. This priority effect arises from iron depletion: the established colonizer depletes free iron to levels that prevent the invader from crossing its survival threshold (**Figure S2**). Thus, despite the presence of distinct siderophore

types, competition for iron as the sole limiting resource enforces the competitive exclusion principle in all four regimes discussed above.

(4) Coexistence: when Eq.(S51) is satisfied, it automatically disables Eq. (S52)-(S53), leaving only two stable state: either both species survive together, or extinct together. It means that coexistence and single-species exclusion are mutually incompatible outcomes: When one species attempts to dominate, the other species, which benefits more from the abundant "foreign" siderophores, will experience a growth surge that restores balance.

### Section 5: Oscillation and Chaos in Three- and Five-Species Systems

#### Modeling the Rock-Paper-Scissor Interactions

In a three-species system (Fig. 3A), each partial-producer secretes one siderophore type but can utilize all three. The resulting Benefit Transfer Graph contains two overlapping "rock–paper–scissors" loops: a counterclockwise loop (species 1 → 2 → 3 → 1) and a clockwise loop (species 3 → 2 → 1 → 3). For simplicity, we set equal self-receptor fractions ($v_{\text{self}}$) across species and parameterize the two cheating loops with $v_{321}$ ($v_{32} = v_{21} = v_{13} = v_{321}$) and $v_{123}$ ($v_{12} = v_{23} = v_{31} = v_{123}$) respectively, with constraint that $v_{\text{self}} + v_{321} + v_{123} = 1$.

#### Attractor Type Assessment

For the three- and five-species siderophore interaction networks (Fig. 2, parameters in Table S2), each parameter set was integrated until the trajectory reached a long-term attractor. Integration was performed automatically by the custom function, which continues simulation in large time blocks until one of the following conditions is satisfied (maximum 5 blocks):

- Steady state: $|dY/dt| < 10^{-3}$ and variance of all species in the second half of the trajectory < 10.
- Limit-cycle: spectral analysis of the trajectory detects a clear dominant frequency with dominant power ratio > 0.3, normalized spectral entropy < 0.3, and harmonic deviation < $10^{-3}$.

Upon convergence, the program returns the final trajectory segment and the time-averaged concentration of each species (mean value over the last periods for limit cycles or over the final stable segment for steady states). The function also reports the detected period (for limit cycles) and flags the attractor type (steady state or limit cycle). Trajectories that failed to satisfy either criterion within the maximum integration blocks were classified as "non-converged/dynamic" (potentially chaotic or complex dynamics) and were grouped with oscillatory cases under the "dynamic" category.
For large scale simulations, the same pipelines were used to perform attractor type assessment.

#### Bifurcation Analysis of the Three-Species System

As self-reliance ($v_{\text{self}}$) increases, the system transitions from oscillatory coexistence to competitive exclusion (no more than one species can survive) (**Figure S3**A). Coexistence is steady when the two cheating loops are balanced ($v_{321} \approx v_{123}$) but becomes oscillatory as imbalances between the loops increase (**Figure S3**B). Overall, coexistence is possible when cheating loops are favored over self-loops in benefit transfer. When coexistence occurs, it is oscillatory if the two cheating loops are sufficiently asymmetric, whereas balanced loops favor steady coexistence (**Figure S3**C)

##### Confirmation of the Heteroclinic Bifurcation

We noticed that for the bifurcation induced by $v_{\mathrm{self}}$ increase, the limit-cycle oscillations exhibit a progressive increase in both amplitude and period (the oscillations becoming markedly slower). Near $v_{\mathrm{self}} \approx 0.28$, the oscillatory behavior abruptly terminates, and the system transitions to a stable steady state (**Figure S4**A). In the context of nonlinear dynamics, this characteristic sequence is the classic signature of a heteroclinic crisis (heteroclinic-mediated global bifurcation) in nonlinear dynamic systems[10].

To elucidate the geometric mechanism terminating the oscillations, we first mapped the system's stationary points via a solution landscape and monitored the minimum Euclidean distance between the stable limit cycles and filtered index-1 saddle points. We observed that as the control parameter asymptotically approaches the critical threshold, the limit cycle expands and moves progressively closer to three specific saddle points (**Figure S4**B-C). This observation indicates that the saddle points “collide” with the limit cycle, effectively breaking the orbital continuity and causing the oscillation period to diverge. Based on the asymptotic deformation of the trajectory and the distances between all the saddle points and the trajectory approaching zero, we conclude that the oscillatory regime is terminated via a heteroclinic bifurcation.

This bifurcation is significant in highlighting abrupt and potentially irreversible transitions in complex systems, where small parameter perturbations can lead to the sudden loss of oscillatory behavior, emphasizing the role of global phase space structures in system resilience. Ecologically, it underscores the tipping points that may result in the collapse of biodiversity-maintaining oscillations into non-coexisting steady states. Such bifurcations have been commonly observed in ecological models incorporating the Allee effect, where reduced population growth at low densities amplifies the risk of overexploitation and sudden regime shifts to extinction or low-diversity equilibria[5].

#### Bifurcation Analysis of the Five-Species Chaotic System

In a more complex five-species system (Fig. 2J), increasing self-reliance ($v_{\mathrm{self}}$) progressively suppresses complexity: chaotic oscillations give way to stable periodic cycles and, eventually, to exclusion (**Figure S5**).

### Section 6: Large Community Simulation

#### Generation and Simulation of Randomized Communities

In simulating large-scale communities, we choose a system with $N_{\text{spe}} = 50$ species that can choose from 50 different siderophores (analogous to the 49 types of lock-key siderophore-receptors pairs in the Pseudomonas iron interaction network [6]).

We characterized cheating behavior along two complementary axes (Fig. 4A): (1) cheating breadth ($CB$), defined as the average number of siderophore types a species can utilize in addition to its own ($\frac{1}{N_{\text{spe}}}\sum_{i=1}^{N_{\text{spe}}}\sum_{j=1}^{N_{\text{sid}}}(v_{ij} > 0\ \wedge \alpha_{ij} = 0)$); and (2) pure-cheater ratio ($r_c$), defined as the percentage of species that produce no siderophores ($\alpha_{i,j} = 0 \text{ for all } j > 0$).

Under given cheating breadth $CB$ and ratio of pure-cheaters $r_c$, the community interaction parameters $\alpha_{ij}$ and $v_{ij}$ are generated by Uniform distribution, where species are designated as producers or cheaters with probability $1 - r_c$ and $r_c$, respectively. Cheaters have growth budget of $\alpha_{i0} = 1$. Producers randomly select a siderophore type $j$ and allocate a production budget $\alpha_{ij}$ drawn uniformly from (0,1): for simplicity, we assumed each species produces at most one siderophore type. $CB$ cheating receptors are assigned randomly across non-self positions in the $v_{ij}$ matrix, and receptor weights are sampled uniformly and normalized. Each different set of $\alpha_{ij}$ and $v_{ij}$ represents a distinct community.

Starting from randomized initial conditions, we simulated each community to its asymptotic state and recorded the number of surviving species, where "surviving" was defined as maintaining a time-averaged biomass greater than $10 \cdot \sigma_i/d$. Such simulations were performed $1.5 \cdot 10^6$ times.

Given the distribution of final biomass (**Figure S6**), we define species maintaining a time-averaged biomass greater than $10^{-3}$ as "high-abundance species".

Chemostat parameters were described in Table S2.

#### Cheating Breadth Non-Monotonically Influences Dynamics and Diversity

We observed dynamic behavior in non-extinct communities followed a non-monotonic pattern (**Figure S7**A): as cheating breadth increased from 0 to 15, the proportion of communities exhibiting oscillatory or chaotic dynamics rose from near 0% to a peak of 5–

11% (depending on pure-cheater ratio from high to low). Diversity is calculated by the Shannon entropy of the time-average biomass of all species in the community [11] (**Figure S7**B). It also exhibits moderate levels of non-monotonicity, especially under high pure-cheater ratio.

#### Ranking of the Benefit Transfer Edges in Community

We analyzed the Benefit Transfer Graph for each non-extinct community. For every producer $i$, we ranked all its outgoing benefit edges ($b_{ij}$) by magnitude (Fig. 4E), starting from 1. These Rank-1 edges represent the strongest benefit flow in this community.

For subgraph formed by all high-abundance species in the community, we calculated the number of in-going Rank-1 edges, then scale it by the number of high-abundance species (Fig. 4F). The enrichment pattern is robust across community states: rank-1 edges constitute the internal structure of the high-abundance subgraph in over 90% of exclusion cases (single survivors), approximately 60% of steady coexistence, and around 80% of dynamic coexistence. This suggests that successful species distinguish themselves by being the high-priority beneficiaries in the BTG, forming densely connected interaction cores of the community. In each community, random species were selected by the same number of high-abundance species, and their subgraphs in BTG does not exhibit any enrichment for high-ranking edges (**Figure S8**).

### Section 7: Core of the Maximal Benefit Transfer Graph Predicts Community Fate

#### Definition of Maximal Benefit Transfer Graph

The set of all Rank-1 benefit-transfer edges forms a reduced subgraph of the BTG, which we termed the maximal Benefit Transfer Graph (mBTG). In mBTG, each node represents a microbial species, and each directed edge indicates the maximal benefit flow from a siderophore producer to the species that derives the greatest growth benefit from it (the maximal beneficiary, **MB**). Thus, every producer node has exactly one outgoing edge, while some species may receive benefits from multiple producers.
For each producer $i$, its maximal beneficiary $j^*$is defined as:

$$j^* = \arg\max_j b_{ij}, \tag{S54}$$

where $b_{ij}$ denotes the growth benefit species $j$ receives from the siderophore produced by species $i$, as defined by Eq. (S8) in the main text.

Thus, each producing species has one outgoing edge, and species that do not produce siderophores (pure-cheater) have no outgoing edges. Species that are maximal beneficiaries (i.e., receive at least one maximal benefit) can have one or more incoming edges.
Due to this "outdegree can only be one or zero" property, The mBTG is a directed pseudoforest, a graph structure defined by the constraint that every node has an out-degree of at most one[12]. It is an important mathematical structure primarily investigated within the fields of graph theory and algorithm design, with practical applications in areas such as cryptography[13] and parallel/distributed computing[14].

Central to a graph are two types of "components," i.e., the "groups of nodes" connected by different standards. A Weakly Connected Component (WCC) is a maximal set of nodes that are connected when all edges are treated as undirected. In contrast, a Strongly Connected Component (SCC) respects edge direction. An SCC is a maximal set of nodes such that every node can reach every other node via directed paths. That is, for any two nodes in an SCC, there is a directed path from one to the other and another path back.

#### Definition of Core in WCC, and Proof for its Existence and Uniqueness

Within this framework, we define the "Core" of a WCC as its Terminal Strongly Connected Component (TSCC)[15]. Formally, it is defined as a minimal non-empty subset of nodes $C$ within a WCC such that no edges leave the set $C$ (i.e., for any edge from $i$ to node $j$, if $i$ is in C, then $j$ is also in $C$). Examples are shown in Fig. 4B.

#### Classification of Cores by Loop Length

We characterize each Core by its Loop Length ($\mathrm{L}$), defined as the number of edges in the subgraph induced by the Core's nodes. This metric rigorously distinguishes the three topological architectures identified in the main text:

1. **Sink Core** ($L = 0$): A TSCC consisting of a single node with an out-degree of 0 (a pure-cheater). This subgraph has 1 node and 0 edges (W0 in Fig. 4B).
2. **Self-loop Core** ($L = 1$): A TSCC consisting of a single node with a self-directed loop (a producer benefiting maximally from itself). The induced subgraph has 1 node and 1 edge (W1 in Fig. 4B).
3. **Cyclic Core** ($L \geq 2$): A TSCC consisting of $k$ nodes ($k \geq 2$) forming a simple cycle. The induced subgraph has $k$ nodes and $k$ edges (W2 and W3+ in Fig. 4B).

#### Existence and Uniqueness of the Core

Every WCC in mBTG contains exactly one Core. The proof follows from the finite nature of the graph and the out-degree constraint ($d_{out} \leq 1$).

1. **Existence**: Consider a path formed by traversing edges starting from any arbitrary node in a finite WCC. Since the graph is finite, the path must eventually either (a) terminate at a node with no outgoing edges (a Sink), or (b) revisit a node, thereby closing a cycle.
   a) In case (a), the terminal node forms a TSCC of size 1 with no edges ($L = 0$).
   b) In case (b), once the path enters a cycle with $n$ nodes, it cannot escape because the nodes in the cycle already utilize their single allowed outgoing edge to point to the next node in the loop. Thus, the cycle forms a TSCC with $\mathrm{L} = n$. $n = 1$ for Self-Loop Core, $n \geq 1$ for Cyclic Core.

   In both cases, a Core exists.

2. **Uniqueness:** Suppose, for the sake of contradiction, that a single WCC contains two distinct Cores, $C1$ and $C2$. Since they belong to the same WCC, there must exist an undirected path connecting a node in $C1$ to a node in $C2$. Because both $C1$ and $C2$ are terminal (no edges leave them), any path connecting them must essentially "flow" into them. Therefore, at some point along the undirected path between $C1$ and $C2$, there must exist a "branching" node $v$ that has directed paths leading to both $C1$ and $C2$. This would require node $v$ (or a downstream node) to have an out-degree of at least 2. This contradicts the definition of the mBTG, where $d_{out} \leq 1$.

**Conclusion:** Consequently, each WCC possesses exactly one structural Core, acting as the unique attractor for all ecological flows within that component.

#### WCCs Maps to Systems' Attractors

In simulations of large communities with $N_{\mathrm{spe}} = 50$ and $N_{\mathrm{sid}} = 50$, we found that each WCCs in the mBTG maps directly to one of the systems' attractors, i.e. "community fate": Regardless of the initial conditions, the final biomass always concentrated entirely within a single WCC rather than distributing across several (**Figure S9**A) . This implies that species belonging to different WCCs effectively occupy separate basins of attraction and seldom coexist. Moreover, biomass within a surviving WCC is highly concentrated in its core (**Figure S9**B): In steady-state communities, nearly all biomass resided in the core; in dynamic communities, the core species contained nearly 50% of the biomass, while some other species fluctuate outside it in the same WCC. Thus, each WCC dictates a specific community fate, and its core serves as the structural and dynamic nucleus of the community.

#### Core Loops Length Predicts Community Fates

We then link the topological properties of the core to specific community fate. In simulation, we generated $3 \cdot 10^5$ random communities, located one of its WCCs, then distributed the biomass randomly to nodes within this WCC. We observed that Core's loop length predicts the final outcome of the community simulation:

1. The probability of global extinction: 97% for WCCs with Sink Cores ($L = 0$); 2% for these with Self-loop Cores ($L = 1$); and increase to a steady value around 20% for these with Cyclic Cores ($L \geq 2$) (**Figure S10**).
2. The probability of entering an steady-state attractor is around 90% for Cores with loop length smaller than 3. When Cores contained three or more species (loop length ≥ 3), oscillatory dynamics became a common outcome (35–40%) (Fig. 4C).
3. In steady-state systems, species richness increased with loop length up to three before plateauing. In dynamic systems, richness continued to rise with loop length (Fig. 4D).

Taken together, each WCC corresponds to a distinct attractor, and its core structure predicts the ecological fate of the community that converges upon it. We made the following simplified prediction rules, for communities with initial biomasses concentrated in one WCC (Examples shown in **Figure S11**):

1. Sink Core ($L = 0$): global extinction (97% accuracy);
2. Self-loop Core ($L = 1$): survival of the single core species (95% accuracy).

3. Cyclic Core of two or more species ($L \geq 2$): coexistence of at least two species in the WCC (69% accuracy). This lower value is due to the approximately 20% extinction rate and the around 10% of cases where the final biomass will concentrate to another more competitive WCC.

### Section 8: Graph Analysis of the Maximal Benefit Transfer Graph

#### Classification of Node Types in mBTG

Graphically, the nodes in the mBTG can be classified into two broad categories: Maximal Beneficiaries (MBs), and Leaf Nodes (Fig. 4E):

**1. Maximal Beneficiaries (MBs)**

A Maximal Beneficiary (MB) is a species $m$ that receives the largest benefit from at least one species, i.e.:

$$\text{exist } i \text{ , such that } m = \arg\max_{j} b_{i,j}., \tag{S55}$$

Based on their siderophore production patterns ($\alpha_{ij}$), MBs can be further divided into three subtypes:

**(a) Self-looped MB**

A species $m$ that is the maximal beneficiary of its own siderophore production, defined as：

$$m = \arg\max_{j} b_{m,j}., \tag{S56}$$

Graphically, its outgoing edge points to itself (a self-loop, length = 1). Biologically, this corresponds to a pure-producer or partial-producer that benefits most strongly from its own siderophore.

**(b) Connector MB**

A maximal beneficiary $m$ that produces siderophore, but does not gain maximal benefit from its own siderophore:

$$\text{exist } i, \text{such that } m = \arg\max_{k} b_{ik}, ;\ and\ \text{ for } j = \arg\max_{k} b_{m,k} , j \neq m. \tag{S57}$$

Graphically, it has one outgoing edge pointing to another MB, and at least one incoming edge. Biologically, it can only be a partial-producer that both produces its own siderophore and effectively exploits others' siderophores. These nodes connect different benefit paths, allowing resource flow between species producing different siderophores.

**(c) Terminator MB**

A species $m$ that does not produce any siderophore ($\alpha_{ij} = 0 \text{ for any } j > 0$) but is a maximal beneficiary of at least one producer.

$$\alpha_{m0} = 1;\ and\ \text{ for } i = \arg\max_{j} b_{m,j} , j \neq m. \tag{S58}$$

Graphically, it has incoming edges but no outgoing edge. Biologically, it is a pure-cheater, receiving but not returning benefits. These nodes represent benefit sinks, terminating the flow of siderophore-mediated interactions within a connected subgraph (WCC).

**2. Leaf Nodes**

A species $m$ represents a leaf node, if it is not the maximal beneficiary of any siderophore:

$$b_{i,m} < \max_k b_{ik} \text{ for all } i. \tag{S59}$$

Leaf nodes therefore have no incoming edges in the graph.

Depending on their production behavior, they can be either producer-leafs ($\alpha_{ij} > 0$) acting as benefit sources for others but do not benefit maximally from any siderophore, or isolated cheaters ($\alpha_{ij} = 0$) neither produce siderophores nor receive maximal benefit, appearing as isolated nodes in mBTGs.

In total, a mBTG thus contains four basic node types: (1) Self-looped MBs as self-benefiting producers; (2) Connector MBs as cross-benefiting partial-producers linking species related to different types of siderophores into one WCC; (3) Terminator MBs that are pure-cheaters acting as benefit sinks in a WCC; (4) Leaf Nodes that are producers or cheaters not central to any maximal benefit flow.

### Increased Cheating Breadth Reduces Self-Looped MB in the mBTG.

The decrease in the number of **self-looped** Maximal Beneficiaries (MBs) with increasing cheating breadth (Fig. 4E) can be understood analytically.

In a community of $N_{\text{spe}}$ species choosing from $N_{\text{sid}}$ siderophores, given pure-cheater ratio $r_c$ and cheating breadth $CB$, there are: (i) $N_{\text{spe}} \cdot (1 - r_c)$ producers in total, and (ii) $N_{\text{spe}} \cdot CB$ cheating receptors distributed across the whole community.

#### Expected Number of Produced Siderophores

Because each producer randomly selects from the $N_{\text{sid}}$ siderophores types, the expected number of siderophores that are actually produced is:

$$S = N_{\text{sid}} \cdot \left(1 - \left(\frac{N_{\text{sid}} - 1}{N_{\text{sid}}}\right)^{N_{\text{spe}} \cdot (1 - r_c)}\right). \tag{S60}$$

For the case $N_{\text{sid}} = 50$ and $N_{\text{spe}} = 50$ and $r_c = 0$, this yields approximately 32 types of siderophores produced in a typical community.

#### Case 1: Zero Cheating Breadth

When the cheating breadth is zero, meaning no species can exploit siderophores produced by others, each producer synthesizes and exclusively utilizes its own siderophore. In this case, for a community with $S$ siderophore types being produced, there are exactly $S$ maximal beneficiaries, and all of them are self-looped MBs, since no partial-producers or pure-cheaters exist when cheating breadth is zero.

##### Case 2: Increasing Cheating Breadth

As cheating breadth increases, species gain additional receptors allowing them to exploit siderophores from other species. For any siderophore type, the average number of producers synthesizing it (contributing self-receptors) is $N_{\text{spe}} \cdot (1 - r_c)/S$, while the average number of species cheating on it (contributing cheating receptors) is $N_{\text{spe}} \cdot CB/S$. Assuming that any recipient has an equal chance of becoming the maximal beneficiary, the probability that one of its producers gets selected is:

$$P_{\text{self-looped}} = \frac{N_{\text{spe}} \cdot (1 - r_c)/S}{N_{\text{spe}} \cdot (1 - r_c)/S + N_{\text{spe}} \cdot CB/S} = \frac{1 - r_c}{1 - r_c + CB}. \quad \text{(S61)}$$

This probability declines hyperbolically as cheating breadth $CB$ increases, implying that self-looped MBs become progressively rarer as more cross-species exploitation occurs (**Figure S12**).

From a graph-theory perspective, self-looped MBs contribute no connectivity to the mBTG, because their outgoing edge returns to themselves. Despite the average out-degree kept to be $1 - r_c$, decreased percentage of Self-Looped MB increases the effective out-degree (i.e. out-degree excluding self-loops). This strengthens connectivity and promotes the formation of multi-node WCCs.

The out-degree distribution (excluding self-loops) becomes:

$$P(d_{out} = 0) = r_c;\ P(d_{out} = 1) = (1 - r_c) \cdot (\frac{CB}{1 - r_c + CB}) \quad \text{(S62)}$$

Thus, increasing cheating breadth raises the probability that a producer's maximal-beneficiary edge targets another species rather than itself.

Each self-looped MB forms the core of a WCC. Simulations show that within such components all other species other than the self-looped MB are competitively excluded (Fig. 4D). Therefore, reducing the number of self-looped MBs is a structural prerequisite for enabling multi-species coexistence.

As cheating breadth increases and self-looped MBs are replaced, WCCs can enlarge and begin supporting stable or dynamic coexistence.

#### Increased Cheating Breadth Biases MB Selection Toward Larger $\alpha_{i0}$.

As cheating breadth increases, the number of Leaf nodes in the mBTG also increases (Fig. 4E, last panel), indicating that the number of Maximal Beneficiaries (MBs) decreases. Meanwhile, each type of siderophore has one MB. This reflects a "concentration" effect: the MB of one siderophore increasingly becomes the MB of multiple siderophores. Correspondingly, the in-degree distribution becomes more dispersed with larger cheating breadth: both the fraction of species with zero in-degree and the probability of exceptionally large in-degree increase (**Figure S13**).

We also observe that the number of Terminator MBs rises with cheating breadth (Fig. 4E, third panel). Such cheater nodes form the core of WCCs that typically collapse into extinction in simulations.

Both phenomena arise from a single underlying mechanism: The probability of a species $i$ being selected as a maximal beneficiary, $P_{MB}(i)$, is positively related to its growth allocation $\alpha_{i0}$, and this bias becomes dramatically stronger with increasing cheating breadth $CB$.

We first formalize this mechanism by studying an equivalent probability problem:

#### Probability Question Setup

Start with an array of $N$ independent random numbers $a_i$, each uniformly distributed between 0 and 1. Randomly choose $K$ of them (without replacement). For each chosen element, draw an independent random multiplier $v_i$ also from uniform distribution from 0 to 1, and compute the product

$$b_i = a_i v_i., i = 1 \ldots K.$$

Among these $K$ products, the largest value is chosen.

The question is: For an element whose original value is $a \in (0,1)$, what is the probability that it will ultimately be selected as the largest product ($P_{\text{win}|a}$)?

#### Step 1. Decompose the Problem into Two Random Processes

For a given element in $a$ to be selected:

1. It must be first selected among the $K$ samples.

$$P(\text{selected}) = \frac{K}{N}. \quad \text{(S63)}$$

2. Conditional on being selected, it must beat the other $K-1$ elements after the random scaling by $v_i$. Thus,

$$P_{\text{win}|a} = \frac{K}{N}\, p(a). \quad \text{(S64)}$$

where $p(a)$ is the probability that this element wins, given its value is $a$.

#### Step 2. Probability of Beating Other Competitors

Let $V \sim \text{Uniform}(0,1)$ be the random multiplier for this element. For a single competitor with value $A_j \sim \text{Uniform}(0,1)$ and multiplier $V_j \sim \text{Uniform}(0,1)$, the probability that a competitor not wining is:

$$\Pr\left(A_j V_j < aV \mid V = v\right) = \int_0^1 \Pr\left(A_j < av/V_j\right) dV_j. \quad \text{(S65)}$$

If we define $b = aV$, we can get the expression

$$q(b) = b - b\ln b, \text{ for } b \in (0,1). \quad \text{(S66)}$$

That's the probability that a single competitor's product is smaller than $aV$.

Since there are $K-1$ independent competitors, given $V=v$,

$$P(\text{win} \mid a, v) = [q(av)]^{K-1}. \tag{S67}$$

Finally, we average over all possible $v \in [0,1]$:

$$p(a) = \int_0^1 [q(av)]^{K-1}\, dv = \int_0^1 [av(1-\ln(av))]^{K-1} dv. \tag{S68}$$

Therefore, given Equation (S64), the overall probability that an element with value $a$ is ultimately selected is

$$P_{\text{win}|a} = \frac{K}{N} \int_0^1 [av(1-\ln(av))]^{K-1} dv. \tag{S69}$$

#### Step 3. Applying the Result to Maximal Beneficiaries in the mBTG

In each community, there are $N_{\text{spe}}$ species in total. That represents $N$ in Equation (S69). For each given siderophores, the expected self-producer number is $N_{\text{spe}} \cdot (1-r_c)/S$, the average cheating receptor number is $N_{\text{spe}} \cdot CB/S$, so in total there are $\frac{N_{\text{spe}}}{S} \cdot (CB + 1 - r_c)$ recipients (Fig. 4F). That represents $K$ in Equation (S69).
Therefore, the probability of a species $i$ becomes the MB for this siderophore is:

$$P_{MB}(\alpha_{i0}) = \frac{CB+1-r_c}{S} \int_0^1 [\alpha_{i0} v(1-\ln(\alpha_{i0} v))]^{\frac{N_{\text{spe}}}{S}(CB+1-r_c)-1} dv. \tag{S70}$$

This function increases monotonically with $\alpha_{i,0}$, and its steepness increases with cheating breadth $CB$. (Fig. 4G).

A useful approximation captures the scaling: $P_{MB}(\alpha_{i0}) \sim \alpha_{i0}^{\frac{N_{spe}}{S}(CB+1-r_c)}$.
Thus, when $CB$ is small, the exponent is small, $P_{MB}$ exhibiting weak dependence on $\alpha_{i0}$; When $CB$ is large, $P_{MB}$ exhibits very strong bias toward large $\alpha_{i0}$. Under large cheating breadth, species with small $\alpha_{i0}$ are almost never selected as MB, while species investing more on their growth become dominant MBs that accumulate multiple incoming edges. Pure-cheaters, as their $\alpha_{i,0} = 1$, become particularly dominating.
This increasing bias explains the emergence of Terminator MBs and the dispersion in the in-degree distribution.

#### Estimate the In-degree Distribution in mBTG

For a species $i$, having an edge in the mBTG pointing from species $j$ to $i$ requires:
(1) species $j$ is a producer (probability $1-r_c$);
(2) species $j$ produces siderophore type $k$ (probability $1/S$), and species $i$ is the maximal beneficiary of that siderophore (probability $P_{MB}$ in Equation (S70)).
Summing over siderophore types cancels the $1/S$ factor, resulting in $P(\text{edge } j \to i) \simeq (1-r_c)\, P_{MB}(\alpha_{i,0})$.
Assuming independence across producers (a first-order approximation), the in-degree of

species $i$ follows a binomial form of $(1-r_c)P_{MB}(\alpha_{i0})$:

$$p_{\text{indeg}}(d,\alpha_{i0}) = \binom{N_{\text{spe}}}{d}\left[(1-r_c)P_{MB}(\alpha_{i0})\right]^{d}\left[1-(1-r_c)P_{MB}(\alpha_{i0})\right]^{N_{\text{spe}}-d}$$

Integrating over the distribution of $\alpha_{i,0}$ gives the community-wide in-degree distribution:

$$P_{\text{indeg}}(d) = \int_0^1 \binom{N_{\text{spe}}}{d}\left[(1-r_c)P_{MB}(\alpha)\right]^{d}\left[1-(1-r_c)P_{MB}(\alpha)\right]^{N_{\text{spe}}-d} d\alpha. \quad \text{(S71)}$$

#### Removing Self-loops to Obtain the Effective In-Degree Distribution

However, self-directed edges do not contribute to connectivity, and Equation (S71) does not exclude self-loops. The probability of a self-loop is given by Equation (S61), Therefore, the effective in-degree (excluding self-loops) is:

$$P_{\text{effectivyIN}}(d) = \begin{cases} P_{\text{indeg}}(d)\left(1-P_{\text{self-looped}}\right) + P_{\text{indeg}}(d+1)P_{\text{self-looped}}, \text{for } d>0 \\ P_{\text{indeg}}(d) + P_{\text{indeg}}(d+1)P_{\text{self-looped}}, \text{for } d=0 \end{cases}. \quad \text{(S72)}$$

Under large $CB$, this distribution becomes highly dispersed, producing both many zero-in-degree nodes and many very-high-in-degree nodes. The theoretical prediction matches simulated networks closely (**Figure S14**), confirming that the observed dispersion arises from the increasing MB-selection bias toward large $\alpha_{i0}$.

### Graph-Theoretic Implications

Dispersed degree distributions are known to promote network percolation, where disconnected components merge into large, connected clusters. We calculated the critical degree of percolation of this system, treating the graph as undirected network (**Figure S15**A) and directed network (**Figure S15**B) respectively. For weakly connected components (WCCs), percolation begins as early as CB>1. For strongly connected components (SCCs), the properties of directed pseudoforest (out-degree ≤ 1) making SCC percolation never fully occurs, though it approaches the threshold.

### Supplementary Figures

Figure S1. Bistability in the Single-Species System. 35
Figure S2. Competitive Exclusion in the Two-Species System. 36
Figure S3. Bifurcation and Phase-Diagram in the Three-Species Oscillation System. 37
Figure S4. Heteroclinic Bifurcation Analysis 38
Figure S5. Bifurcation in the Five-Species Chaos System. 40
Figure S6. Biomass Distribution for the Large Community Simulation 41
Figure S7. How Cheating Breadth Influences Dynamics and Diversity in Large Communities 42
Figure S8. Normalized Frequency of BTG Edge Ranking, for Randomly Drawn Species 43
Figure S9. Biomass Distribution across WCC and Their Cores 44
Figure S10. Probability of Community Extinction as a Function of Core Loop Length 45
Figure S11. Examples of WCCs in a Community and their Corresponding Fate 46
Figure S12. Effects of Cheating Breadth on Self-looped MB and Effective Out-degree 47
Figure S13. Cheating Breadth Increases the Dispersion of In-degree Distributions in the mBTG 48
Figure S14. Estimation of the Effective In-degree Distribution 49
Figure S15. Cheating Breadth Decreases Percolation Thresholds in the mBTG by Reshaping Degree Distributions. 50

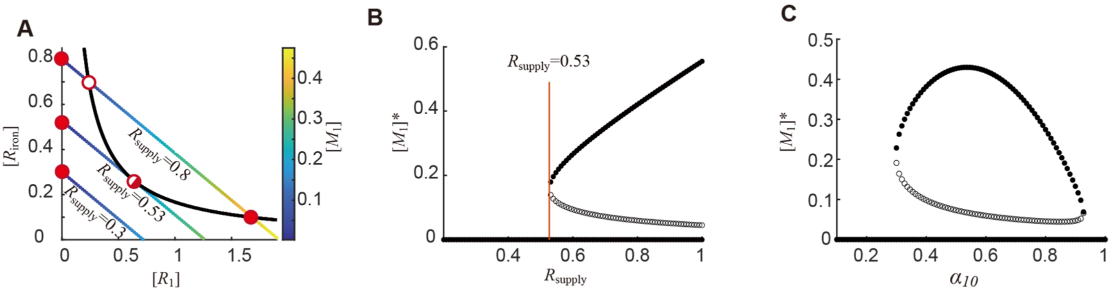

**Figure S1. Bistability in the Single-Species System.**

A. The chemical space of the system, showing the growth contour (black curve) and flux-balance lines for three different values of $R_{\text{supply}}$, represented by colored straight lines. The color indicates the steady-state biomass of the species. Stable fixed points are marked by filled red circles, the unstable fixed point by an empty red circle, and the critical bifurcation point by a half-red, half-white circle.

B. Bifurcation diagram corresponding to the system in (A), illustrating the change in fixed points as $R_{\text{supply}}$ varies. Filled circles represent stable steady-state biomass values, while empty circles indicate unstable fixed points. The threshold value of $R_{\text{supply}}$ calculated using Equation (S22), is indicated by the orange line.

C. Bifurcation diagram as in (B), but showing the system's response to variations in $\alpha_{1,0}$.

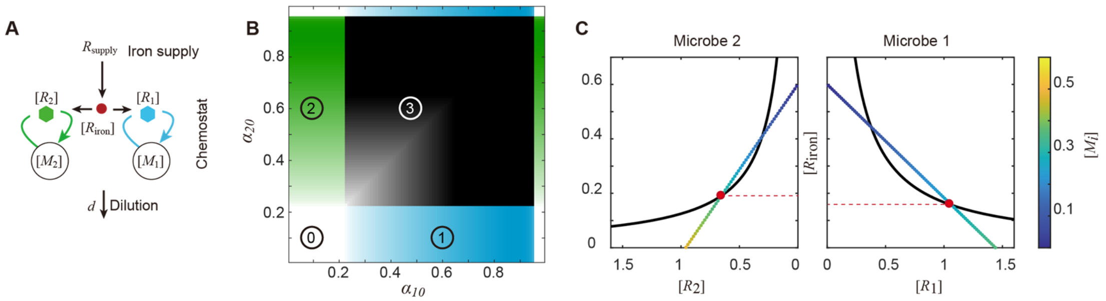

**Figure S2. Competitive Exclusion in the Two-Species System**.

A. Schematic of a two-species system where two pure-producers compete. Each species secretes and exclusively utilizes its own siderophore type (blue for species 1, green for species 2).

B. Phase diagram spanning the growth-allocation space ($\alpha_{1,0}$-$\alpha_{2,0}$) for the competitive system in (A), with fixed $v_{11} = v_{22} = 1$. Distinct ecological outcomes are color-coded and numbered, as demonstrated in Fig. 2G: (0) Global extinction (white); (1) Dominance species 1 and exclusion of species 2(blue); (2) Dominance species 2 and exclusion of species 1 (green); and (3) Priority effect (black). Color intensity is proportional to the total steady-state biomass.

C. The chemical spaces of species 2 (left panel) and species 1 (right panel). For each species, the growth contour is shown as a black curve, and the flux-balance line is represented by colored straight line, with color indicating the steady-state biomass of each species. The red markers indicate the steady-state iron concentrations shaped by the two species in the system.

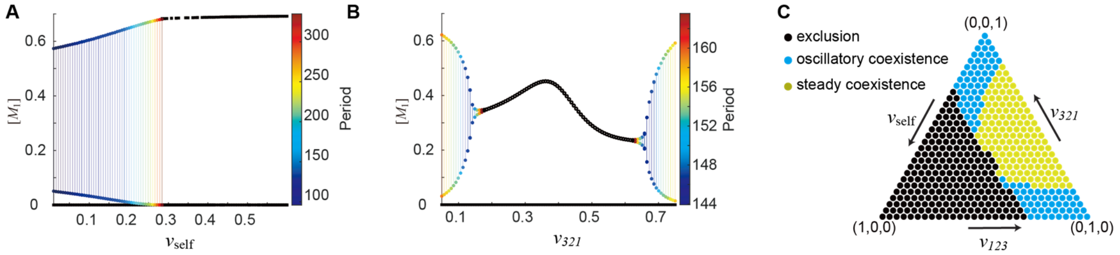

**Figure S3. Bifurcation and Phase-Diagram in the Three-Species Oscillation System**.

A. Bifurcation diagram illustrates transitions from oscillatory coexistence to exclusion as $v_{\mathrm{self}}$ increases.

B. Bifurcation diagram show transitions from oscillatory to steady coexistence then back to oscillation, as $v_{321}$ increases.

C. Phase diagram of system dynamics in the $v_{\mathrm{self}}$-$v_{123}$-$v_{321}$ ternary space, showing regions of exclusion (black), oscillatory coexistence (blue), and steady coexistence (yellow).

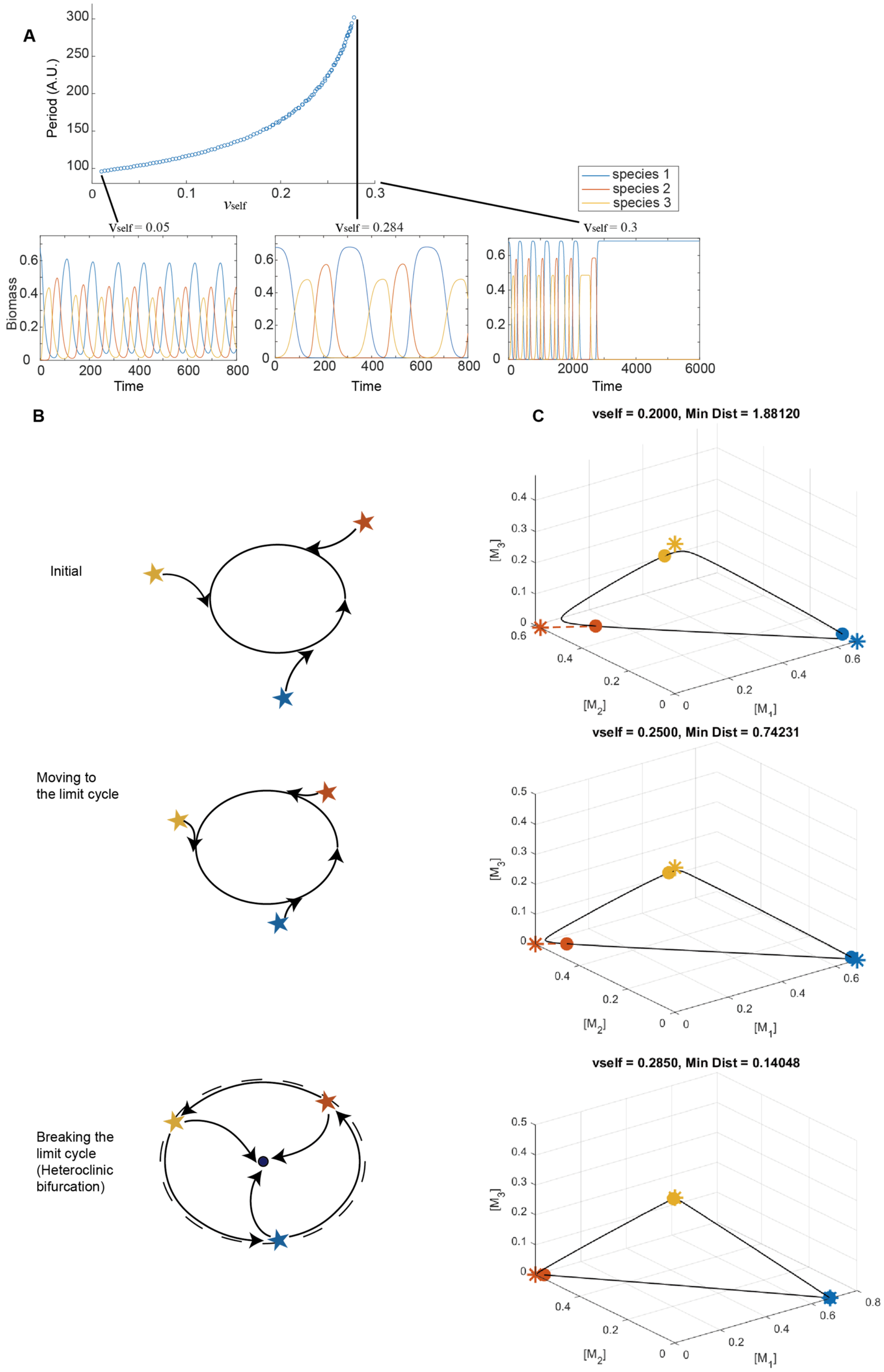

**Figure S4. Heteroclinic Bifurcation Analysis**

A. Top panel shows the monotonic increase in oscillation amplitude as the self-receptor weight $v_{\mathrm{self}}$ is increased. Bottom three panels display representative time series of species biomass at $v_{\mathrm{self}} = 0.05, 0.284, 0.3$, respectively. For low $v_{\mathrm{self}}$, the system exhibits fast limit-cycle oscillations. As $v_{\mathrm{self}}$ increases, the oscillatio period grow, suggesting critical slowing down. After $v_{\mathrm{self}}$ exceeding the bifurcation point, oscillations abruptly collapse and the system transitions to a stable steady state, consistent with a heteroclinic crisis. This global bifurcation marks a sudden loss of cyclic coexistence and a shift to non-oscillatory dynamics.

B. The schematic diagram of heteroclinic bifurcation. The ellipse stands for a limit cycle, the blue node means an attractor and the stars means the index-1 saddles.

C. The phase diagram (projected to the biomass space) at different $v_{self}$. The stars stand for the index-1 saddle points and the cycles stands for the nearest points to the corresponding saddle points in the limit cycle. The blue line stands for the limit cycle.

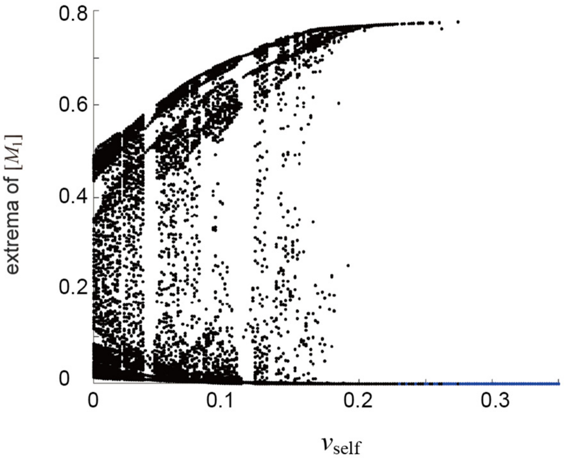

**Figure S5. Bifurcation in the Five-Species Chaos System**.
Bifurcation diagram showing transitions from chaos to exclusion as the self-receptor fraction ($v_{\mathrm{self}}$) increases.

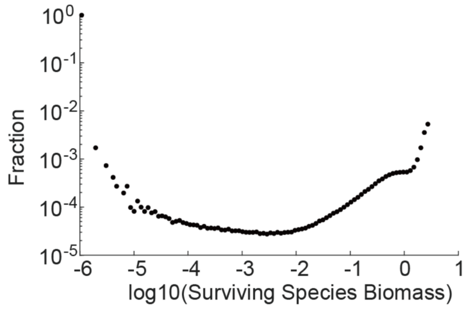

**Figure S6. Biomass Distribution for the Large Community Simulation**

The biomass distribution of surviving species (maintaining a time-averaged biomass greater than $10 \cdot \sigma_i / d$) in log-log scale.

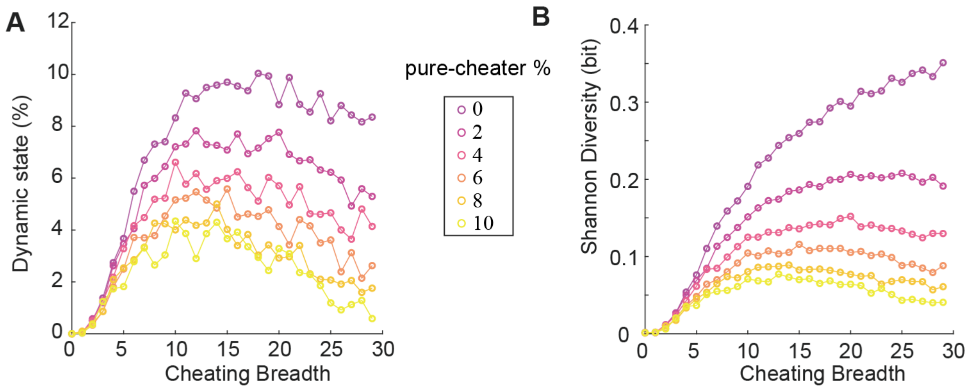

**Figure S7. How Cheating Breadth Influences Dynamics and Diversity in Large Communities**

A. In surviving communities, the probability of exhibiting dynamic states (oscillations or chaos) peaks at intermediate cheating breadths.

B. Average alpha diversity of the community generally increases with the cheating breadth, but exhibits moderate levels of non-monotonicity under high pure-cheater ratios.

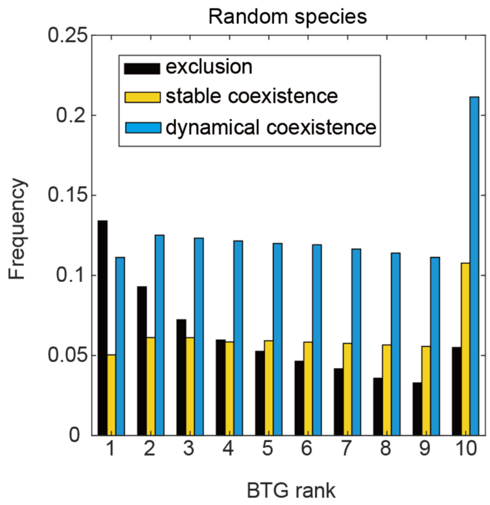

**Figure S8. Normalized Frequency of BTG Edge Ranking, for Randomly Drawn Species**

Rank frequency distribution of benefit transfer edges in BTGs, for random subset species drawn from the community.

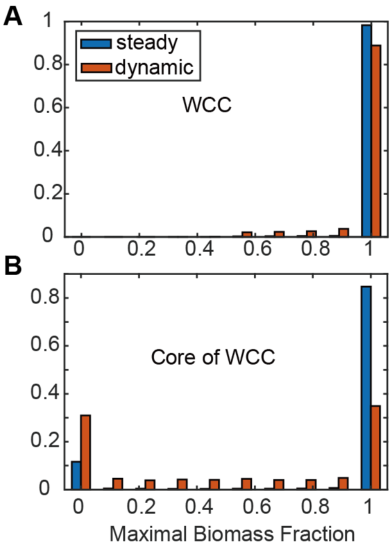

**Figure S9. Biomass Distribution across WCC and Their Cores**

(A) For all non-extinct simulations, the maximal fraction of total community biomass located within each WCC. Biomass overwhelmingly concentrates on a single WCC.

(B) Fraction of total biomass located specifically to the core of the dominant WCC. Even in dynamic communities, the core retains nearly half of total biomass.

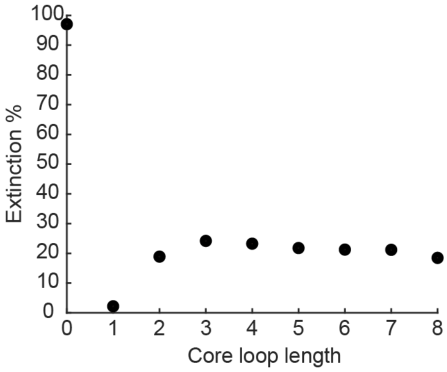

**Figure S10. Probability of Community Extinction as a Function of Core Loop Length**
Simulations are performed from initial conditions distributed to one of the WCC in the community, and the WCC's Core loop length influences the probability of community-level extinction.

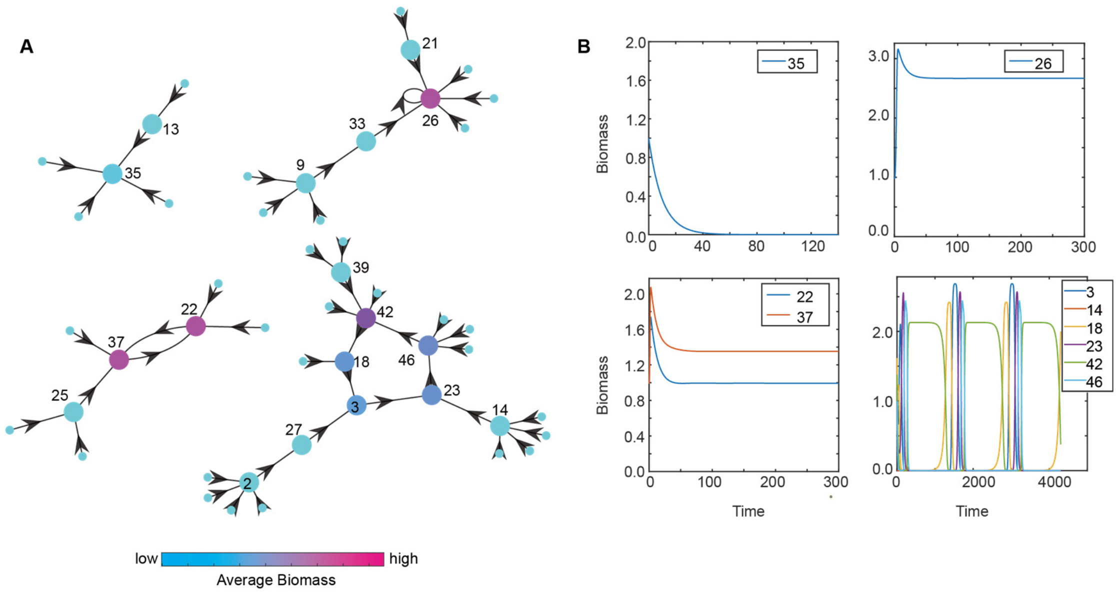

**Figure S11. Examples of WCCs in a Community and their Corresponding Fate**

(A) Example mBTG containing four WCCs with different core types. Node labels denote Maximal Beneficiary species' ID; small pale nodes indicate Leafs without incoming edges. Node colors represent average biomass in asymptotic states across many simulations with randomized initial conditions.

(B) Representative biomass trajectories corresponding to the four WCC cores shown in (A), illustrating four possible fates: extinction, exclusion, steady coexistence, and oscillatory coexistence.

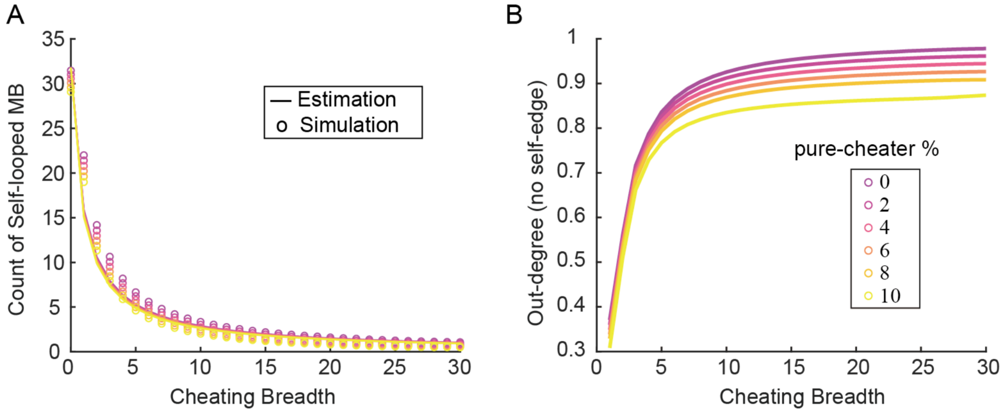

**Figure S12. Effects of Cheating Breadth on Self-looped MB and Effective Out-degree**

(A) The number of Self-looped MB decreases hyperbolically as cheating breadth increases, consistent with the analytic prediction in Eq. (S61).

(B) Effective out-degree (excluding self-loops) increases with cheating breadth. Larger cheating breadth reduces self-loops and increases the likelihood that maximal-beneficiary edges point to other species.

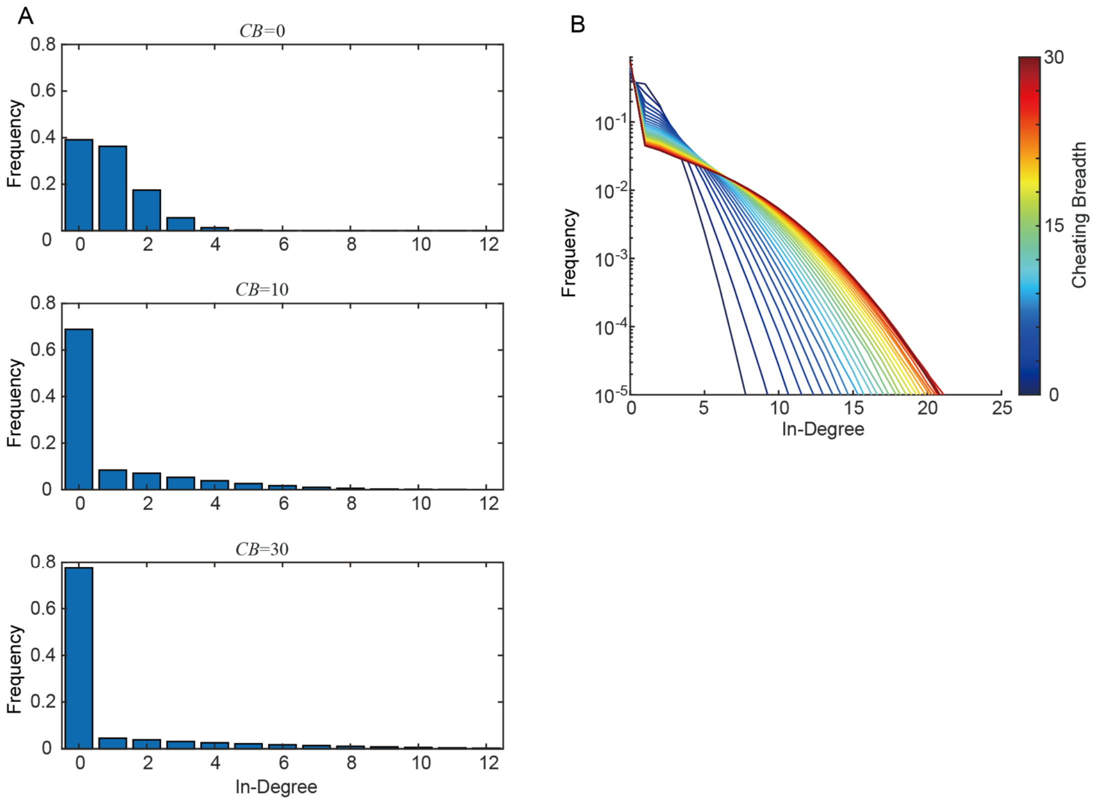

**Figure S13. Cheating Breadth Increases the Dispersion of In-degree Distributions in the mBTG**

(A) Empirical in-degree histograms of the maximal-beneficiary graph (mBTG) at low (CB = 0), intermediate (CB = 10), and high (CB = 30) cheating breadth. Increasing CB suppresses moderate in-degrees while inflating both zero-degree and high-degree frequencies.

(B) In-degree distributions shown across cheating breadth values. Larger CB produces increasingly heavy-tailed distributions.

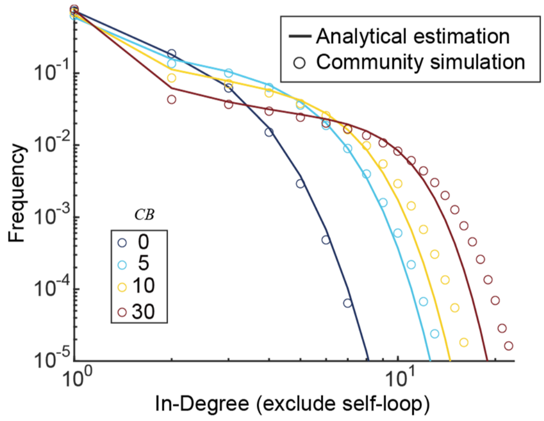

**Figure S14**. **Estimation of the Effective In-degree Distribution**

Analytical estimation is obtained by Eq. (S72), and shown by curve. Community simulation results are shown by circles.

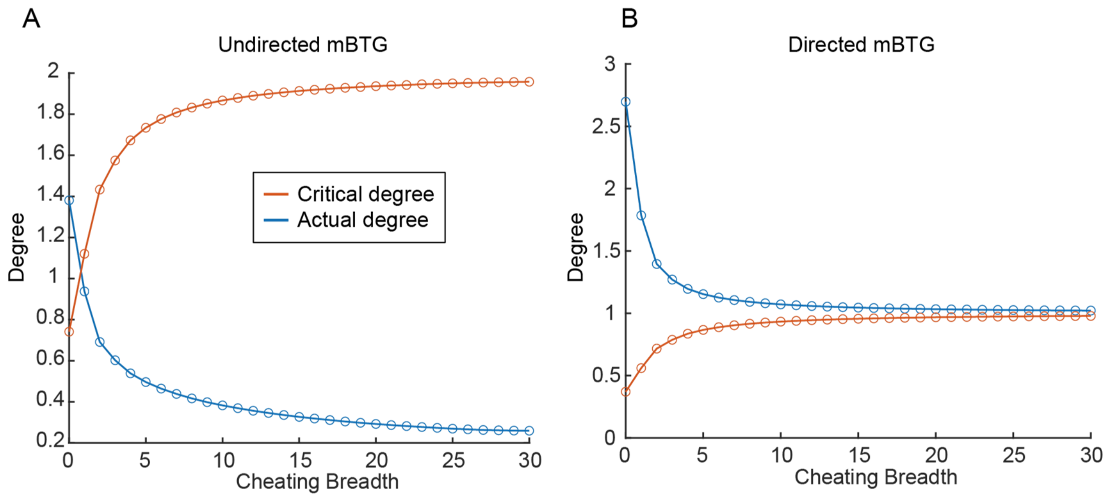

**Figure S15. Cheating Breadth Decreases Percolation Thresholds in the mBTG by Reshaping Degree Distributions.**

(A) Undirected percolation analysis. The effective out-degree of the system (excluding self-loops) is obtained by simulating $10^6$ graphs (Blue curve). Treating the mBTG(mBTG) as an undirected network, the average degree crosses the classical percolation threshold is shown in red curve.

(B) Directed percolation analysis. Same as (A), other than that mBTG is treated as a directed network.
